# Non-identifiability and the Blessings of Misspecification in Models of Molecular Fitness

**DOI:** 10.1101/2022.01.29.478324

**Authors:** Eli N. Weinstein, Alan N. Amin, Jonathan Frazer, Debora S. Marks

## Abstract

Understanding the consequences of mutation for molecular fitness and function is a fundamental problem in biology. Recently, generative probabilistic models have emerged as a powerful tool for estimating fitness from evolutionary sequence data, with accuracy sufficient to predict both laboratory measurements of function and disease risk in humans, and to design novel functional proteins. Existing techniques rest on an assumed relationship between density estimation and fitness estimation, a relationship that we interrogate in this article. We prove that fitness is not identifiable from observational sequence data alone, placing fundamental limits on our ability to disentangle fitness landscapes from phylogenetic history. We show on real datasets that perfect density estimation in the limit of infinite data would, with high confidence, result in poor fitness estimation; current models perform accurate fitness estimation because of, not despite, misspecification. Our results challenge the conventional wisdom that bigger models trained on bigger datasets will inevitably lead to better fitness estimation, and suggest novel estimation strategies going forward.

## 1 Introduction

The past decades have witnessed a tremendous increase in the scale of genome sequence data available from across life. Recently, methods for estimating molecular fitness using generative sequence models have seen widespread success at translating this evolutionary data into predictions of the functional consequences of mutation. Such models have been shown to accurately predict the outcomes of experimental assays of protein function [23, 44, 36], and have been applied to infer 3D structures of RNA and protein [35, 56] and to design novel proteins [50, 47, 33]. The models have also been used to predict whether human mutations are pathogenic, directly informing the diagnosis of genetic disease [19]. In this paper, we investigate how and why generative sequence models fit to evolutionary sequence data are successful at estimating molecular fitness, and how they might be improved and generalized going forward.

Existing approaches to fitness estimation with generative sequence models rest on an assumed relationship between density estimation and fitness estimation. Given a dataset of sequences *X*_1_, …, *X*_*N*_, assumed to be drawn i.i.d. from some underlying distribution *p*_0_, fitness models proceed by (1) fitting a probabilistic model *q*_*θ*_ to *X*_1:*N*_ and (2) using the inferred density 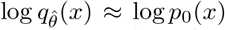 as an estimate of the fitness *f* (*x*) of a sequence *x*; this estimate in turn is used to predict other covariates such as whether the mutated sequence is pathogenic [23, 44, 19]. Innovation in fitness models has come out of a trend of building increasingly flexible models fit to increasing amounts of data: simple models that treat each column of a sequence alignment independently were improved by energy-based models that accounted for epistasis [23], which in turn were improved by deep variational autoencoders [44], which in turn were improved by deep autoregressive alignment-free models [50, 33, 36]. Naively, one might assume that these improvements have come from obtaining better and better estimates of the data distribution *p*_0_, and improvements will continue with bigger models and bigger datasets. In this article, we argue that this presumption is incorrect.

### Technical summary

First, we show that that the true data distribution *p*_0_ may not reflect fitness, and argue instead that we should be focused on estimating another distribution that does, *p*^∞^ (the “stationary distribution”, to be defined below). In particular, we demonstrate that phylogenetic effects – i.e. the history of how current sequences evolved over time – can “distort” the observed data, leading to a situation where *p*_0_ ≠ *p*^∞^ (Sec. 2). Second, we show in this situation that *p*^∞^ and fitness *f* are non-identifiable: even with infinite data, there always exists some alternative fitness function 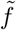 that explains the same data just as well as *f*. This sets fundamental limits on what we can learn about fitness from evolutionary data (Sec. 3). Third, although exact estimation of *p*^∞^ is impossible, we show that it is still possible to get closer to *p*^∞^ than *p*_0_, that is, to find a better estimator of fitness than the true data density *p*_0_. This can be done by fitting to data a parametric generative sequence model ℳ = {*q*_*θ*_ : *θ* ∈Θ} that is (approximately) well-specified with respect to *p*^∞^ (i.e. *p*^∞^∈ ℳ) but *misspecified* with respect to the data distribution *p*_0_ (i.e. *p*_0_ ∉ */*ℳ), thus illustrating how misspecification can be a blessing rather than a curse (Sec. 4). Fourth, we construct a hypothesis test to determine whether the blessings of misspecification occur on real data, for existing fitness estimation models; our test uses a recently developed Bayesian nonparametric sequence model to construct a credible set for *p*_0_ (Sec. 6). Fifth, we apply our test to over 100 separate sequence datasets and fitness estimation tasks, to conclude that existing fitness estimation models systematically outperform the true data distribution *p*_0_ at estimating fitness (Sec. 7). The takeaway is that better fitness estimation (i.e. better *p*^∞^ estimation) will not come from better density estimation (i.e. better *p*_0_ estimation); bigger models and bigger datasets are not enough. Instead, better fitness estimation can come from developing models that describe *p*^∞^ better but the data density *p*_0_ *worse*.

## 2 Models of Fitness and Phylogeny

In this section we illustrate how *p*_0_ may not accurately reflect the true fitness landscape, by analyzing a generative model of sequence evolution that takes into account not only fitness but also phylogeny. Our model is general: it allows for arbitrarily complex epistatic fitness landscapes, and recovers standard phylogenetic and fitness models as special cases. Our concerns about the effects of phylogeny on fitness estimation are motivated by the widespread use – and trust – of phylogenetic models for evolutionary sequence data (phylogenetic models are far more widely applied than fitness models) [21, 12, 17, 18]. Note, however, that phylogeny is only one reason why *p*_0_ may not not reflect the true fitness landscape, and that our later results on the benefits of misspecification in fitness models (Sec. 4) do not depend on the specific cause of mismatches between *p*_0_ and the fitness landscape.

### Joint fitness and phylogeny models

We define “joint fitness and phylogeny models (JFPMs)” using two elements: a description of how individual species (or populations or individuals) change over time, which depends on fitness *f*, and a description of the species’ relationship to one another, a phylogeny **H**. To describe the dynamics of individual species, let *P* ^*τ*^ (*x, x*_0_) denote the probability of sequence *x*_0_ evolving into sequence *x* after time *τ* ; in particular, *P* ^*τ*^ (*x, x*_0_) is assumed to be the transition probability of an irreducible continuous-time Markov chain defined over sequence space 𝒳. For example, under neutral evolution (i.e. without selection based on fitness), *P* ^*τ*^ (*x, x*_0_) may follow a Jukes-Cantor model [18]. With selection, for simple population genetics models (e.g. Moran or Wright processes), Sella and Hirsh [49] demonstrate under general conditions that for any *x*_0_,

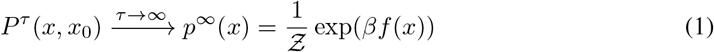

where *f* (*x*) is the log fitness of the sequence *x* and *β* > 0 is a constant (Appx. B). The implication of Eqn. 1 is that the stationary distribution of the evolutionary dynamics follows a Boltzmann distribution, with energy proportional to the log fitness of the sequence. Estimating *p*^∞^ is of interest because it provides a direct estimate of log fitness, up to a linear transform, since *f* (*x*) = *β*^−1^(log *p*^∞^(*x*) + log *Ƶ*). (N.b. in the remainder of the paper, when we say “estimate fitness” we mean, implicitly, “estimate log fitness up to a linear transform”.)

The sequences we observe, however, do not necessarily come from the stationary distribution. Instead, they are correlated with one another according to their evolutionary history. This is described by a phylogeny **H** = (*V, E, T*) consisting of a directed and rooted full binary tree with edges *E* and nodes *V*, along with time labels for the nodes, *T* : *V* → ℝ_+_ (Fig. 1A). Each node *v* is associated with a sequence *X*_*v*_, drawn as 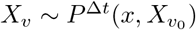, where 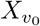 is the sequence of the parent node, *v* is the child node, and Δ*t* = *T* (*v*_0_) −*T* (*v*) is the length of the edge between them (Fig. 1B). The root sequence is drawn from *p*^∞^. The observed datapoints *X*_1_, …, *X*_*N*_ correspond to the leaf nodes. In general we will assume all leaves are observed at effectively the same time, the present day *T* = 0.

**Figure 1:**
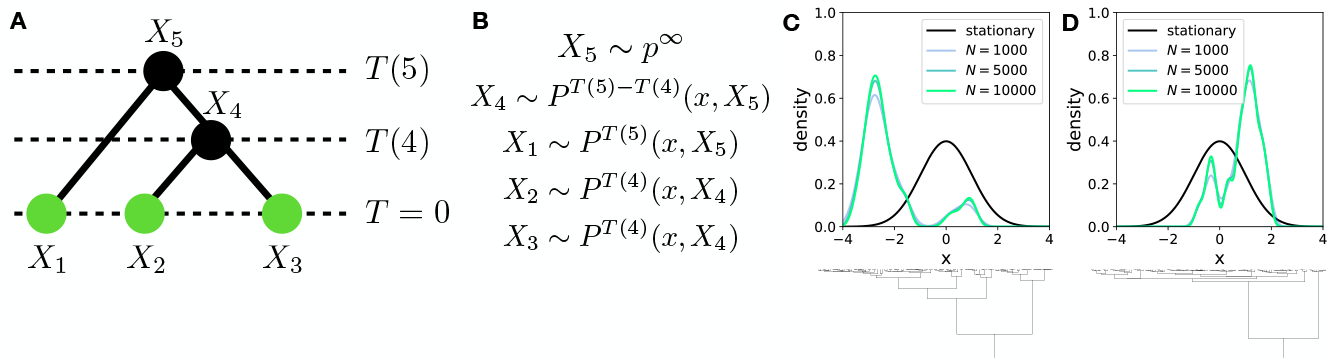
JFPM illustration. (A) Example JFPM for *N* = 3 observed sequences. (B) Generative process for sequences at each node of the phylogeny **H**. (C) *Above:* Stationary distribution *p*^∞^ and kernel density estimates of the distribution of samples *p*_0_ from an OUT model for increasing *N*. *Below:* A subset of the phylogeny. (D) Same as (C) for an independent sample of **H**.

### Special cases

Standard probabilistic phylogenetic models ignore fitness and assume

#### Assumption 2.1

(Pure phylogeny models (PMs)). *Constant fitness: f* (*x*) = *C*.

Example models that fit this form include most of those used in BEAST [14], MrBayes [25], RaxML [51], etc. Standard probabilistic fitness models, on the other hand, ignore phylogenetic history and assume that the stationary distribution has been reached,

#### Assumption 2.2

(Pure fitness models (FMs)). *Let τ*_*i*_ *be the distance in time between observed sequence X*_*i*_ *and its parent node. Take τ*_*i*_ → ∞ *for all i, which implies that*

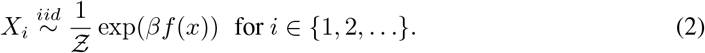

The key implication of this assumption is that density estimation and fitness estimation are linked: the data follows *X*_1_, …, *X*_*N*_ ∼ _*iid*_ *p*_0_ = *p*^∞^, and so if we can estimate *p*_0_ we can estimate the fitness. Example models include EVMutation [23], DeepSequence [44], EVE [19], etc. Note although Assumptions 2.1 and 2.2 do not conflict directly, conclusions made based on them conflict in practice: PMs typically infer finite and different lengths for branches (i.e. *τ*_*i*_ *<* ∞), while FMs typically infer differences in fitness (i.e. *f* (*x*) ≠ *C*), even when applied to the same dataset.

**1D Example** If Asm. 2.2 does *not* hold, then there is no reason for the distribution of observed sequences *X*_1_, *X*_2_, … to follow *p*^∞^. We illustrate this with an example, the most widely used JFPM that does not use Assumptions 2.1 or 2.2: the Ornstein-Uhlenbeck tree (OUT) model [18, 8]. In this model, *X* is continuous, i.e. *X* ∈ ℝ, and evolves on a quadratic (fitness landscape of the form) *f* (*x*) ∝ (*x* − *μ*)^2^ +*C* according to the dynamics *P* ^*τ*^ (*x, x*) = Normal 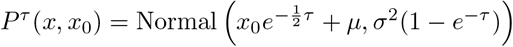. The stationary distribution *p*^∞^ is Normal(*μ, σ*^2^). One can show (Appx. C.1) that for any **H,**

#### Proposition 2.3

(OUT observations). *The distribution of observed genotypes X*_1:*N*_ *is drawn from a multivariate normal distribution with mean* 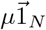 *and covariance* ∑ *where*

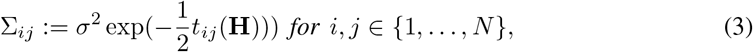

*and t*_*ij*_(**H**) *is the total time of the shortest path between leaves i and j along the phylogeny* **H**.

We drew samples from the OUT with a Kingman coalescent prior on **H** ([3] Def. 2.1) and plotted their density (Fig. 1C). Even as *N* → ∞, the distribution of samples does not follow *p*^∞^. Moreover, rerunning the process with a new sample from the prior yields a very different distribution (Fig. 1D).

## 3 Non-identifiability

In this section we investigate whether, given infinite sequence data, it is possible to infer fitness *f* without Asm. 2.2, and conversely, whether it is possible to infer phylogeny **H** without Asm. 2.1. That is, we are interested in whether fitness and phylogeny are identifiable in general JFPMs. We conclude they are not: given infinite data generated with any *f* and **H**, there exists some alternative 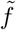 and 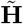, where 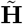 satisfies Asm. 2.2, that explains the data equally well.

Naively, this result may be surprising: in FMs, each sequence is drawn independently, i.e. *X*_*i*_ ⫫*X*_*j*_ |**H**, *f*, while in JFPMs and PMs there is (in general) correlation between sequences, i.e. 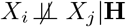, *f*. One might then hope that examining correlations between sequences would enable us to infer whether Asm. 2.2 holds. However, we can show that these correlations are uninformative due to a symmetry in JFPMs, exchangeability.

### Assumption 3.1

(Exchangeability). *Let m*(*X*_1_, *X*_2_, …) *denote the marginal probability of an infinite set of sequences X*_1_, *X*_2_, … *integrating over all phylogenies, i*.*e. m*(*X*_1_, *X*_2_, …) = ∫ *p*(*X*_1_, *X*_2_, … |**H**)*p*(**H**)*d***H**. *Then, for any permutation π of the integers*,

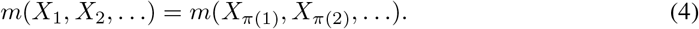

Exchangeability says that if we had observed the sequences in a different order, it would not change their probability. Exchangeability is a ubiquitous assumption in machine learning and statistics models, and its application depends primarily on the information available in a dataset: it is a sensible assumption whenever the ordering of the datapoints provides no useful information. In typical datasets used for fitness estimation, sequences are separated by millions of years of evolution, and are thus all effectively observed at the same time: the present day, *T* = 0. In other words, there is no *a priori* way of ordering the sequences in the dataset, and so we must assume exchangeability. Standard priors on phylogenetic trees, such as the Kingman coalescent, are explicitly constructed to enforce exchangeability [3, 14].

Exchangeability implies that fitness and phylogeny are not identifiable. Even if *X*_1_, *X*_2_, … are generated from a JFPM with a finite branch length phylogeny **H**, we can describe the same data just as well using an FM model with an infinite branch phylogeny 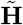 :

### Theorem 3.2

(Non-identifiability). *Assume X*_1_, *X*_2_, … *satisfy Assumption 3*.*1. Then with probability 1 there exists some function* 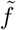 *such that*

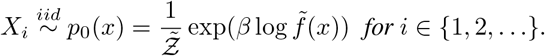

*Proof*. Applying de Finetti’s Theorem ([29], Thm. 11.10), a.s. there exists a random measure *G* such that for *i* ∈ {1, 2, …}, 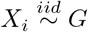. Let *p*_*G*_(*x*) be the pmf of *G*. (We assume *x* is a finite discrete sequence; we can also use continuous *x* assuming the pdf *p*_*G*_(*x*) exists.) Set 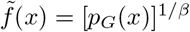. This result says that the observed sequences from an exchangeable JFPM, *X*_1_, *X*_2_, …, are precisely i.i.d. samples from some *p*_0_. Although in the standard tree representation 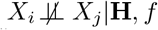, *f*, there must be some alternative description of the same process where 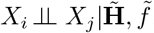. Fitness and phylogeny are thus non-identifiable: data generated from a JFPM with fitness *f* and phylogeny **H** can be described just as well using 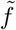 and 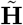, and vice versa. We emphasize that this non-identifiability result is highly general, and does not depend on the specific choice of evolutionary dynamics *P*^*τ*^, only on the assumption of exchangeability.

The biological intuition behind Thm. 3.2 is that if two sequences are similar to each other and distant from a third, they may be similar either because they are closely related (i.e. the distance *τ* to the most recent common ancestor is small) or because they are in a local maximum of the fitness landscape. Without further assumptions, we cannot tell the difference between these two explanations. The machine learning intuition is that evolution, as described by a JFPM, is in effect a Markov chain Monte Carlo process whose stationary distribution gives the fitness. However, the samples we observe may not be fully independent: each pair of samples was initialized from the same point (the most recent common ancestor), and the burn-in since that point may not be sufficiently long. Without independent samples, our estimate of the stationary distribution will be biased.

### Fitness inference as hyperparameter inference

While general, Thm. 3.2 is not constructive, and does not tell us what the distribution *p*_0_ actually is, or how exactly it differs from *p*^∞^. Thm. 3.2 also leaves unclear how much we need to know to learn the fitness landscape: could we infer fitness *f* if we knew the parametric form of *p*^∞^, i.e. if we had some model ℳ and knew that *p*^∞^ ∈ ℳ? What if we also knew the underlying phylogeny **H**? In the long branch limit (Asm. 2.2), fitness is identifiable if **H** is known; if ℳ is also known, learning fitness is a matter of inferring model parameters. In the limit where all the branch lengths in the phylogeny are zero, the distribution of observations from a JFPM reduces to *X*_1_ ∼*p*_∞_ and *X*_1_ = *X*_2_ = *X*_3_ = …. Here fitness is non-identifiable even if **H** and ℳ are known; learning fitness is a matter of learning from a single sample. In the realistic intermediate branch length case, if **H** and ℳ are known, we will show that learning fitness is essentially a matter of *hyperparameter* rather than *parameter* inference.

We demonstrate this last claim in the context of the OUT example, by approximating the OUT model as a Gaussian process latent variable model (GPLVM). We find that fitness only appears as a hyperparameter of the derived GP, not as a parameter. The GPLVM has latent variables *Z*_1_, *Z*_2_, … that lie on the hyperbolic plane ℍ, and uses the Gaussian process kernel *k*(·, ·) = exp(−*d*(·, ·)), where *d*(·, ·) is a distance metric over ℍ. Let 𝒲_1_(·, ·) be the Wasserstein metric for distributions over infinite matrices, i.e. over ℝ^∞×∞^, using the sup norm on matrices.

#### Theorem 3.3

(GPLVM approximation of OUT). *Assume a prior over phylogenies* **H** *that is exchangeable in its leaves and where the minimum time between any pair of nodes is greater than η >* 0 *with probability 1. Define the leaf distance matrix* 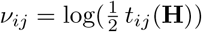. *For any ϵ >* 0, *there exists a*.*s. a GPLVM of the form*,

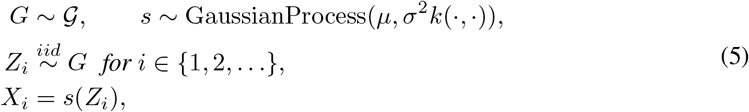

*where G is a random measure over* ℍ, *such that* 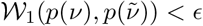, *where* 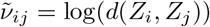.

*If 𝒲*_1_(*p*(*ν*), 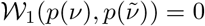, *the OUT and GPLVM produce identical distributions over X*_1_, *X*_2_, … *a*.*e*.. The proof is in Appx. C.2, and uses the embedding of Sarkar [48]. This result says that, by embedding phylogenies **H** in a metric space, we can approximate an OUT arbitrarily well with a GPLVM; as the Wasserstein bound gets smaller, the distribution of covariance matrices of the two models get closer. In the GPLVM, the observations are conditionally independent, *X*_*i*_ ⫫*X*_*j*_ |*s, G*, in line with Thm. 3.2. The phylogeny **H** enters the GPLVM only through the latent space embedding *Z*_1_, *Z*_2_, Learning phylogeny, given the fitness landscape, is thus essentially a matter of inferring latent variables [44, 13]. The fitness landscape enters the GPLVM only through the prior on the Gaussian process (i.e. through *μ* and *σ*). Inferring fitness given phylogeny is thus essentially a matter of inferring hyperparameters. This is both good and bad news for fitness inference. On the one hand, hyperparameters are often learned in practice, and doing so can yield substantially better predictions, so we should be able learn something about *μ* and *σ* given data ([58], Chap. 5). On the other hand, hyperparameters are in general (though not always) non-identifiable, and therefore so is fitness [34]. Ho and Ané [22] describe non-identifiability conditions for the OUT in particular. We conclude that even when **H** andℳ are known, fitness inference in JFPMs is fundamentally challenging.

## 4 Benefits of misspecification

We have demonstrated that a plausible biological mechanism – namely, phylogenetic effects – can produce a data distribution *p*_0_ that does not reflect fitness, and can make exact inference of fitness impossible even given infinite data. Nonetheless, the practical success of fitness estimation methods suggest it is possible to at least approximate fitness landscapes from observational sequence data.

Recall that existing methods proceed by fitting a probabilistic model *q*_*θ*_∈ ℳ = {*q*_*θ*_ : *θ* ∈Θ} to data *X*_1:*N*_, typically via maximum likelihood estimation or approximate Bayesian inference, and then using the predicted log density log 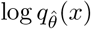 as an estimate of the fitness of a sequence *x*. Why is this approach empirically successful? In this section we consider two hypotheses, either of which may hold true in theory. In Secs. 6-7 we develop and apply tests to evaluate them on real data.

Note that our results in this and the following sections are independent of the specific evolutionary mechanisms that generate a mismatch between *p*_0_ and *p*^∞^, i.e. they are not specific to phylogenetic effects or JFPMs, nor do they even depend on the existence of a stable reproductive fitness function *f* over evolutionary time. We can, in fact, redefine *p*^∞^ to be an arbitrary “target distribution”, with log *p*^∞^ proportional to any chosen measure of molecular fitness or function (such as enzyme activity, fluorescence, etc.). For the sake of concrete illustration, however, we will continue to focus on JFPMs as our primary example of why the data distribution, *p*_0_, may not equal the target distribution we want to estimate, *p*^∞^.

We consider two hypotheses for the empirical success of existing fitness estimation methods.

### Hypothesis #1

(informal). *Fitness estimation methods succeed by finding* 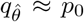, *since for all practical purposes on real data, p*_0_ = *p*^∞^.

This hypothesis would make sense, in JFPMs, if Asm. 2.2 held, i.e. branch lengths were long enough in real datasets for 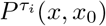 to be close to its stationary distribution. Under this hypothesis, better density estimators have been, and will continue to be, better fitness estimators. We should focus on developing models ℳ that are well-specified with respect to the data, i.e. *p*_0_ ∈ ℳ (Fig. 2A).

**Figure 2:**
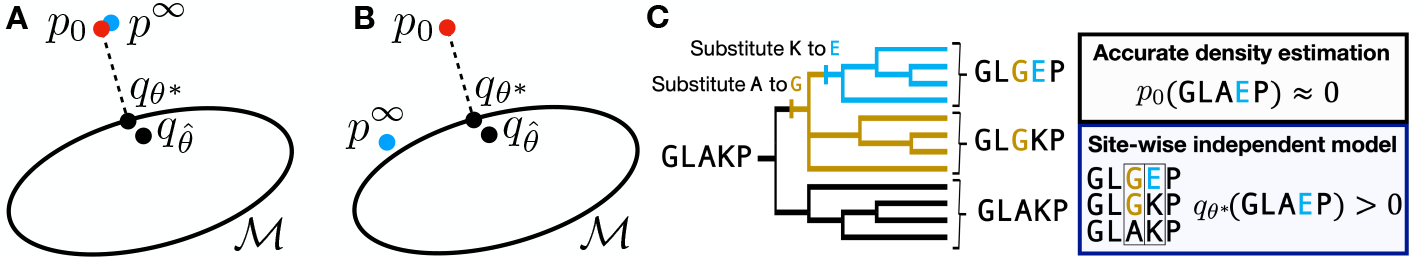
Alternative explanations for the success of fitness estimation methods. (A) Setup in which Hypothesis 1 would hold true. (B) Setup in which Hypothesis 2 would hold true. (C) Biological intuition for the benefits of misspecification (Hypothesis 2).

### Hypothesis #2

(informal). *Fitness estimation methods succeed by using models ℳ that are misspecified with respect to p*_0_, *i*.*e. p*_0_ ∉ ℳ. *The inferred model* 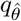 *is then closer to p*^∞^ *than p*_0_ *is*.

To show this second hypothesis is plausible, we prove that it is guaranteed to hold under general conditions. We study the projection of *p*_0_ onto ℳ via the Kullback-Leibler (KL) divergence, 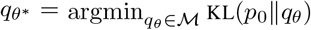. The KL projection is relevant because maximum likelihood estimation minimizes the approximate KL divergence between the data and the model, and the posterior in “log-convex”, meaning that for any *θ, θ*′ ∈ Θ and 0 < *r* < 1, there exists some *θ*^′ ′^ such that Bayesian inference asymptotically concentrates around the maximum likelihood estimator [38]. We thus expect the fit model 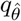 to be close to *q*_*θ*_*, and get closer with *N*. Assume that ℳ is *q*_*θ*_*′′* (*x*) = *q*_*θ*_(*x*)^*r*^*q*_*θ*_*′* (*x*)^1 −*r*^*/*∑ _*x*_ *q*_*θ*_(*x*)^*r*^*q*_*θ*_*′* (*x*)^1 −*r*^; examples of log-convex models include the Potts model, as well as all other exponential family models. Let TV(*p*∥*q*) be the total variation distance between *p* and *q*, and let ∥*g*∥_∞_ = sup_*x*_ |*g*(*x*)| be the uniform (sup) norm of *g*.

### Theorem 4.1

(Benefits of misspecification). *Assume that the model* ℳ *is log-convex and that q*_*θ*_* *exists and is unique. If the model is “less misspecified” with respect to the stationary distribution p*^∞^ *than with respect to the data distribution p*_0_, *in the sense that*

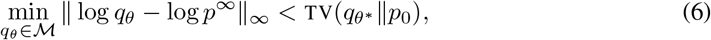

*then*

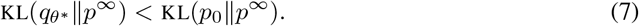

*However, if the model is well-specified with respect to the data distribution, i*.*e. p*_0_ ∈ ℳ, *we have*,

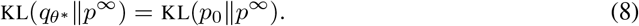

Sec. C.3 contains the proof, and explains how Thm. 4.1 can be extended with more general conditions (we also emphasize again that the proof does not make any assumption that *p*_0_ and *p*^∞^ follow a JFPM). Thm. 4.1 says that if the model is less misspecified with respect to the target distribution than with respect to the data distribution, then projecting the data distribution onto the model will yield a distribution *q*_*θ*_* closer to the target distribution. In other words, the best models for fitness estimation are those at a “sweet spot” of complexity, flexible enough to capture *p*^∞^ but not so flexible as to capture *p*_0_.

To understand the biological intuition behind this result, consider a situation where two neutral mutations with no effect on fitness occur successively at different sites (Fig. 2C). Due to phylogenetic correlation, there is no observed sequence *x*\* in which the second mutation is present but not the first, so an accurate density estimator will find *p*_0_(*x*\*) ≈ 0. However, if we can guess correctly that the fitness landscape is independent across sites, then fitting a site-wise independent model ℳ will imply the mutation is allowed, *q*_*θ*_* (*x*\*) > 0, correctly inferring *p*^∞^(*x*\*) > 0.

Under Hypothesis 2, progress in the field of fitness estimation has *not* come from building better density estimators (Hypothesis 1), but rather from an iterative process of (1) hypothesizing, based partly on biophysical knowledge, models that are approximately well-specified with respect to *p*^∞^ but poorly specified with respect to the data distribution *p*_0_, and then (2) comparing their density estimates against experimental fitness measurements. We will show that on real data, Hypothesis 1 can often be rejected in favor of Hypothesis 2.

## 5 Related Work

Efforts to account for the effects of phylogeny in fitness estimation have a long history [32]. Practical generative sequence models that explicitly account for both epistatic fitness landscapes and phylogeny have long been sought, but stymied primarily by computational challenges [28, 46]. In their place, a variety of non-generative (and often heuristic) methods for correcting for phylogeny have been proposed, including data reweighting schemes [35, 46], data segmentation schemes [10], post-inference parameter adjustments [16], covariance matrix denoising methods [42], simulation based statistical testing [45], and more. In this article, we show that deconvolving fitness and phylogeny is not just computationally hard, but also in general statistically impossible: fitness and phylogeny are non-identifiable. We further show that use of a misspecified parametric model can on its own (without further corrections) partially adjust for phylogenetic effects.

Our results also intersect with the literature on robust statistics: we can think of the observed data distribution *p*_0_ as a “distorted” version of the true distribution of interest *p*^∞^. However, in typical robust inference frameworks (e.g. Huber’s epsilon contamination model), the observed distribution differs from the true distribution by the addition of outliers [24, 52]. Our setup is, in some sense, the opposite: inliers are deleted, as phylogenetic correlations can result in an effective support of *p*_0_ that is *smaller* than that of *p*^∞^ (Fig. 1CD).

## 6 Diagnostic Method

In this section, we develop diagnostic methods to discriminate between Hypothesis 1 and Hypothesis 2 (Sec. 4) based on observational sequence data and experimental fitness measurements, and validate these diagnostics in simulation. Recall that under Hypothesis 2, the estimate 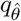 from a parametric fitness model is a better estimate of fitness than the true data density *p*_0_, while under Hypothesis 1, *p*_0_ is better. Discriminating these two hypotheses on real data is nontrivial because we do not have access to *p*_0_. Ideally, then, a diagnostic test would evaluate the probability that the true density *p*_0_ outperforms 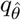 at predicting fitness, taking into account uncertainty in what *p*_0_ could actually be, given the data. To accomplish this, we compute a posterior over *p*_0_ using a Bayesian nonparametric sequence model. In particular, we apply the Bayesian embedded autoregressive (BEAR) model, which can be scaled to terabytes of data and satisfies posterior consistency ([2], Thm. 35):

### Theorem 6.1

(Summary of BEAR posterior consistency). *Assume p*_0_ *is subexponential, i*.*e. for some t >* 0, 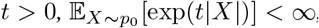, *where* |*X*| *is the length of sequence X. Assume the conditions on the prior detailed in Amin, Weinstein and Marks [2]. If X*_1_, *X*_2_, … ∼ *p*_0_ *i*.*i*.*d*., *then for M >* 0 *sufficiently large and ϵ* ∈ (0, 1*/*2) *sufficiently small*,

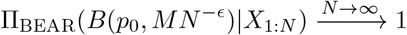

*in probability, where B*(*p, r*) *is a Hellinger ball of radius r centered at p, and* Π_BEAR_(·|*X*_1:*N*_) *is the BEAR posterior*.

Crucially, this result implies that the BEAR posterior will converge to effectively any value of *p*_0_, no matter what *p*_0_ is (unlike a parametric model’s posterior). Moreover, BEAR quantifies uncertainty in its estimates, giving the range of possible values of *p*_0_ that are consistent with the evidence.

We construct our diagnostic test by comparing the fitness estimation performance of 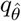 to the range of possible performances of *p*_0_ estimated by BEAR. Let 𝒮_*f*_ (*p*) be a scalar score evaluating how accurately a density *p* predicts fitness *f*. In practice, 𝒮_*f*_ will be based on experimental and clinical measurements of quantities directly related to fitness.

**Diagnostic test** (Test Hypothesis 1 vs. Hypothesis 2.) *Hypothesis 1* 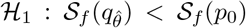. *Hypothesis 2* 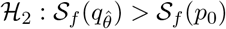. *Accept Hypothesis 2 at significance level α >* 0 *if*

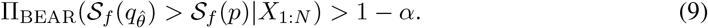

*Accept Hypothesis 1 at significance level α if*

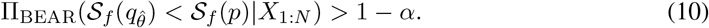

So long as 𝒮_*f*_ (*p*) is a well-behaved function of *p* (in particular, so long as 𝒮_*f*_ is continuous in a neighborhood of *p*_0_ with respect to the topology of convergence in total variation), Thm. 6.1 implies that this diagnostic test will be asymptotically consistent, in the sense that it converges to the correct hypothesis in probability.

### Simulations

We next evaluate the performance of our diagnostic test on simulated data. We considered two scenarios, the first in which Hypothesis 1 holds, and the second in which Hypothesis 2 holds. In both, we let ℳ be a site-wise independent (SWI) model, in which each position of the sequence is drawn independently, i.e. *X*_*l*_ ∼ Categorical(*v*_*l*_) for *l*∈ {1, …, |*X* |}. The parameter *v*_*l*_ is in the simplex Δ_*B*_, where *B* + 1 is the alphabet size. (Further details in Appx. D.) In Scenario 1, the true data are generated according to a Potts model and *p*_0_ = *p*^∞^. In this scenario, the SWI model is misspecified, and misspecification is *bad*: using a more flexible model will produce an asymptotically more accurate estimate of *p*^∞^. We find that our diagnostic test asymptotically correctly accepts Hypothesis 1, in line with Thm. 6.1 (Figs. 3A and 7A). In Scenario 2, the true data are generated according to a JFPM with finite branch lengths, and *p*^∞^∈ ℳ while *p*_0_ ∉ ℳ. The mutational dynamics *P* ^*τ*^ follow the Sella and Hirsh [49] process. The phylogeny ℋ is drawn from a Kingman coalescent. In this scenario, the SWI model is again misspecified, but misspecification is *good*: while the nonparametric BEAR model can achieve better density estimates than the SWI model (Fig. 3C), the SWI model outperforms BEAR at fitness estimation (Figs. 3D and 8). We find that our diagnostic test correctly accepts Hypothesis 2 (Figs. 3B and 7B).

**Figure 3:**
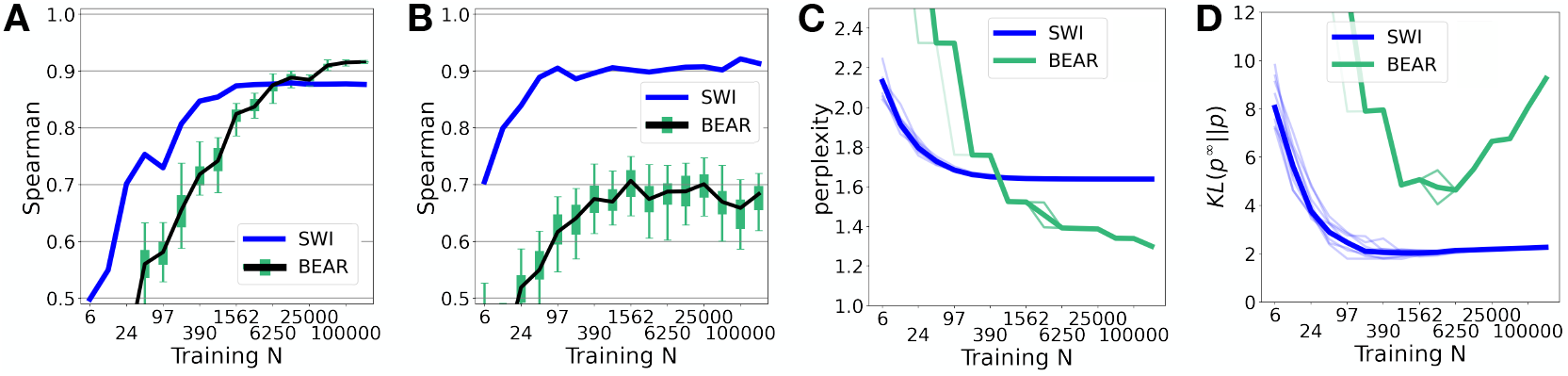
BEAR diagnostic applied to simulated data. (A) Scenario 1. Spearman correlation between the maximum likelihood SWI model and the true fitness 𝒮_*f*_ 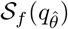, compared to the BEAR posterior distribution over 𝒮_*f*_ (*p*). Quantiles and 95% credible interval shown with green box and whisker. Points above (below) the whiskers correspond to SWI models that significantly outperform (underperform) the true data distribution. (B) Same as A, for Scenario 2. (C) Perplexity on heldout data of the BEAR and the SWI models in Scenario 2. Thick line corresponds to the average over 10 individual simulations (thin lines). (D) Same as C, comparing the KL divergence to *p*^∞^.

A possible point of concern is that the test is poorly calibrated from a frequentist perspective, and in the low *N* regime accepts Hypothesis 2 in Scenario 1 more than 100*α*% of the time when the data is resampled from *p*_0_ (Fig. 9A). This behavior is common in nonparametric Bayesian tests, and not necessarily a problem: the test is still valid from a purely Bayesian perspective. Nevertheless, on real data we will check that we are close to the large *N* regime by (1) checking that the BEAR posterior predictive is at least as close to *p*_0_ as 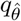 is (as measured by perplexity on held out data; Figs. 3C and 9B) and (2) examining the plot of the BEAR posterior over 𝒮_*f*_ (*p*) as a function of *N* (as in Fig. 3AB), to check that it has converged.

## 7 Empirical Results

We now evaluate whether existing fitness estimation methods outperform the true data density *p*_0_, i.e. whether we can reject Hypothesis 1 in favor of Hypothesis 2 on real data.

### Tasks

We consider two key prediction tasks where fitness models are applied in practice. The first task is to predict whether variants of a protein are functional, according to an experimental assay of protein function; the metric 𝒮_*f*_ (·) is the Spearman correlation between *p*(*x*) and the assay result [23]. There are typically ∼1000s of measurements per assay. The second task is to predict whether a variant of a protein observed in humans causes disease, according to clinical annotations; the metric 𝒮_*f*_ (·) is the area under the ROC curve when *p*(*x*) is used to predict whether or not a variant is pathogenic [19]. There are typically only a handful of labels for each gene. For the first task, we considered 37 different assays across 32 different protein families, and for the second task, 97 genes across 87 protein families; for each protein family, we assembled datasets of evolutionarily related sequences, following previous work. Note that across the 37 assays and 97 genes, the data used for 𝒮 _*f*_ comes from different experiments and different clinical evidence, often collected by different laboratories or doctors. Thus, our overall conclusions should be robust to the choice of 𝒮 _*f*_.

### Models

We considered three existing fitness estimation models: a site-wise independent model (SWI), a Bayesian variational autoencoder (EVE [19], which is similar to DeepSequence [44]), and a deep autoregressive model (Wavenet) [50]. Note that SWI and EVE, unlike Wavenet, require aligned sequences as training data. Details in Appx. E.

### Results

Applied to the first prediction task, our diagnostic test accepts Hypothesis 2 at significance level *α* = 0.025 in 35/37 assays (95%) for SWI, 35/37 assays (95%) for EVE, and 36/37 assays (97%) for Wavenet (Fig. 4A). Applied to the second prediction task, our diagnostic test accepts Hypothesis 2 at significance level *α* = 0.025 in 31/97 genes (32%) for SWI and 46/97 genes (47%) for EVE (Fig. 4B). Thus, fitness estimation models are capable of outperforming the true data distribution *p*_0_. We found evidence for Hypothesis 1 in only a handful of examples: on the first task, Hypothesis 1 was accepted at significance level *α* = 0.025 in 0/37 assays for SWI, 1/37 assays (3%) for EVE, and 0/37 assays for Wavenet, while on the second task, Hypothesis 1 was accepted for 5/97 genes (5%) for SWI and 4/97 genes (4%) for EVE. We confirmed that the diagnostic test was in the large *N* regime: BEAR outperformed Wavenet at density estimation, providing better predictive performance on 27/37 assays (73%) and similar performance on the remaining 10 assays (Fig. 10). (Note that we cannot do this comparison for SWI or EVE since they are alignment-based [57].) Example plots of the BEAR posterior’s convergence with *N* on the first prediction task showed convergence to values of 𝒮 _*f*_ well below that for parametric fitness estimation models (Figs. 4C and 11-12). Overall, we conclude that there is strong evidence that existing fitness estimation methods reliably outperform the true data distribution *p*_0_ across a range of datasets and tasks.

**Figure 4:**
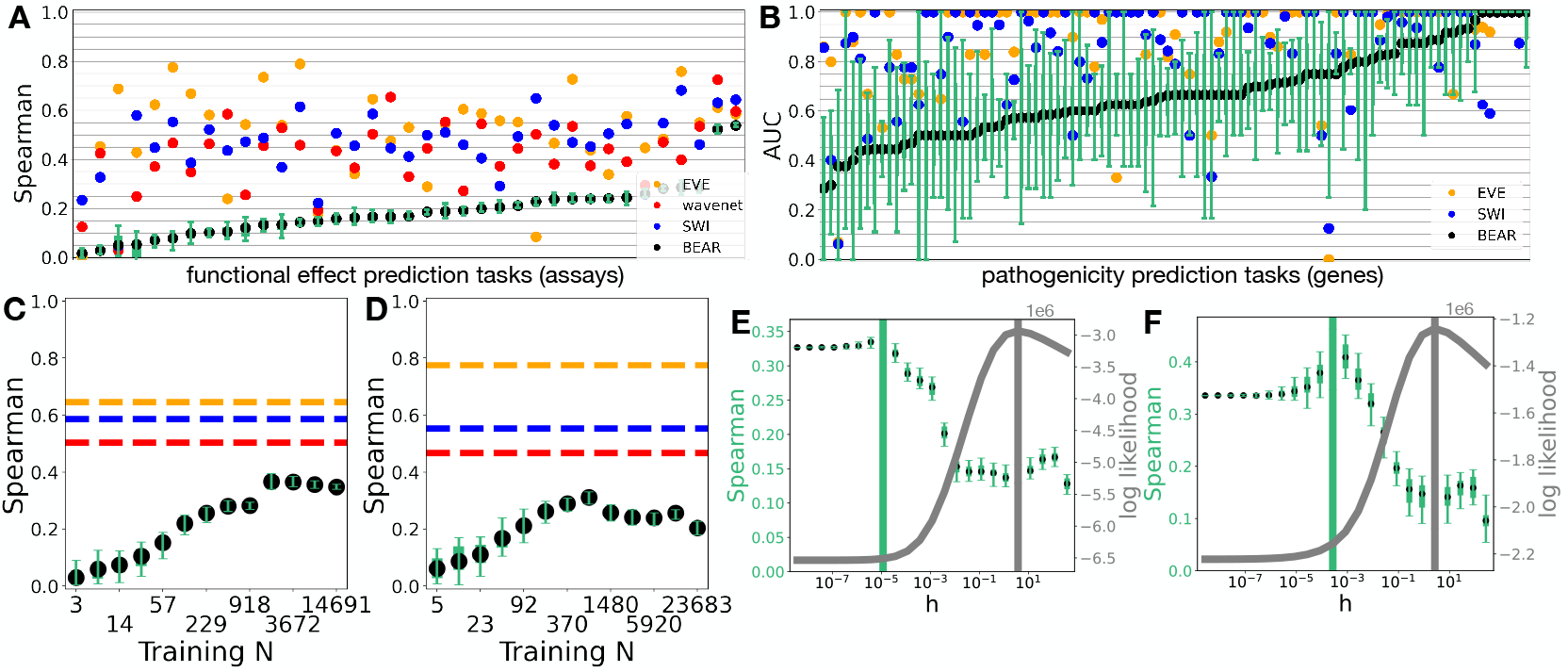
Fitness estimation models systematically outperform the data distribution. (A) Results for the first prediction task, predicting functional measurements in experimental assays. Quantiles and 95% credible interval of the BEAR posterior are shown with the green box and whisker plot. Points above (below) the whiskers correspond to fitness estimation models that significantly outperform (underperform) the true data distribution. (B) Results for the second prediction task, predicting variant pathogenicity in human genes. (C) Convergence of the BEAR posterior with datapoints *N*, for an example assay (*β*-lactamase). (D) Same as C, for another example assay (TIM barrel). (E) BEAR posterior Spearman (black and green) versus BEAR log likelihood (gray), interpolating between parametric and nonparametric regimes (low and high *h*), for an example assay (another *β*-lactamase assay). Peak Spearman indicated with vertical green line, peak log likelihood with gray. (F) Same as E, for another example assay (GAL4 DNA-binding domain).

To study the tradeoffs between density estimation and fitness estimation in more depth, we smoothly and nonparametrically relaxed a parametric autoregressive (AR) model (Appx. E.4). We embedded the AR model (a convolutional neural network) into a BEAR model, and fit the BEAR model with empirical Bayes. We found evidence that the AR model was misspecified on every dataset, following the methodology of Amin, Weinstein and Marks [2]: the optimal *h* selected by empirical Bayes was on the order of 1 − 10 in each dataset. Now, in the limit as the hyperparameter *h* →0, the BEAR model collapses to its embedded AR model; so by scanning *h* from low to high values we can interpolate between the parametric and nonparametric regime. We find a smooth tradeoff between 𝒮_*f*_ (*p*) and the likelihood of the data under the BEAR model, with higher *h* corresponding to better density estimation but worse fitness estimation (Fig. 4EF and 13). This relationship held across many datasets: the diagnostic test, evaluated against the AR model (the *h* →0 limit), accepts Hypothesis 2 in 28/37 assays (76%), but Hypothesis 1 in only 6/37 (16%) (Fig. 14). These results confirm that making a model well-specified (relaxing from a parametric to a nonparametric model) can bring improved density estimation at the cost of worse fitness estimation.

## 8 Discussion

In this article, we have argued that better density estimation does not necessarily lead to better fitness estimation. Further, we estimate with high probability that existing fitness estimation methods systematically outperform the true training data density. Although existing methods rely on flexible, high-parameter deep neural network models, they can nonetheless be misspecified; but this misspecification acts as a blessing, rather than a curse for fitness estimation. Successful models are at a sweet spot of complexity, flexible enough to capture the target fitness distribution well but not so flexible as to match the data distribution itself.

We have focused on state-of-the-art fitness estimation methods which are trained on data from individual protein families [44, 50, 19]. Recently, large-scale generative sequence models (“protein language models”) trained on more diverse datasets (containing proteins from many different families) have show fitness estimation performance comparable to, and in some settings surpassing, single family models [36, 40, 41]. Although applying our diagnostic test to these datasets requires further work, there is no reason to expect the same limitations of density estimation do not hold for such models. Indeed, following a preprint of this paper, Nijkamp et al. [40] presented evidence of the benefits of misspecification in a protein language model: past a certain number of parameters, density estimation improved while fitness estimation deteriorated. See Appx. G for further discussion.

One future direction is to further explore models ℳ that are *less* flexible than existing models and *worse* at density estimation, since they can increase the gap between KL(*q*_*θ*_* ‖ *p*^∞^) and KL(*p*_0_ ‖ *p*^∞^) (Thm. 4.1). There may also be opportunity to improve model geometry: while exponential family models are guaranteed to be log-convex (and thus can satisfy Thm. 4.1), we have no such guarantee for variational autoencoders or other neural network methods. Meanwhile, uncertainty quantification is crucial for applications such as those in clinical genetics, but challenging in misspecified models [53, 39, 26]. Alternatively, it might be useful to abandon the strategy of using misspecified models for fitness estimation altogether, and instead construct JFPM models where the fitness landscape is represented explicitly as a latent variable. Recent progress on amortized variational inference for phylogenetic models is promising for building flexible and scalable JFPMs [55]. However, handling non-identifiability is challenging, and may require new assumptions and/or new methods of sensitivity analysis to infer the full set of fitness landscapes consistent with the data [9].

Although this article has focused on technological applications of fitness models in solving prediction problems, fitness models also have implications for our fundamental understanding of evolution. Pure phylogeny models and pure fitness models present very different pictures of the past history of life: in PMs, similarities and differences among genetic sequences are determined primarily by history and ancestry (Asm. 2.1), while in FMs they are primarily determined by functional constraints (Asm. 2.2). PMs and FMs also present very different implications for the future of life: in PMs, the diversity of sequences seen in nature will likely expand dramatically going forward, while in FMs, the landscape of functional sequences has already been well-explored. Our results emphasize that where and to what extent each model offers an accurate picture of reality remains an open question.

## Acknowledgements

We thank members of the Marks lab, Pascal Notin, Ali Madani, John Ingraham and the anonymous reviewers for insights and suggestions. E.N.W.’s work was supported by the Fannie and John Hertz Foundation. D.S.M. is supported by the Chan Zuckerberg Initiative.

## Checklist

1. For all authors…
  a. Do the main claims made in the abstract and introduction accurately reflect the paper’s contributions and scope? [Yes]
  b. Did you describe the limitations of your work? [Yes] See e.g. Sec. 6 on calibration and Appx. G on evaluating very large models.
  c. Did you discuss any potential negative societal impacts of your work? [Yes] Appx. A
  d. Have you read the ethics review guidelines and ensured that your paper conforms to them? [Yes]
2. If you are including theoretical results…
  a. Did you state the full set of assumptions of all theoretical results? [Yes] Secs. 2 - 6 and Appx. C.
  b. Did you include complete proofs of all theoretical results? [Yes] See Appx. C.
3. If you ran experiments…
  a. Did you include the code, data, and instructions needed to reproduce the main experimental results (either in the supplemental material or as a URL)? [Yes] See supplemental material for code and instructions, and see Appx. E for data.
  b. Did you specify all the training details (e.g., data splits, hyperparameters, how they were chosen)? [Yes] See Appx. E
  c. Did you report error bars (e.g., with respect to the random seed after running experiments multiple times)? [Yes] See Fig. 4.
  d. Did you include the total amount of compute and the type of resources used (e.g., type of GPUs, internal cluster, or cloud provider)? [Yes] See Appx. E.2.
4. If you are using existing assets (e.g., code, data, models) or curating/releasing new assets…
  a. If your work uses existing assets, did you cite the creators? [Yes] See Appx. E.
  b. Did you mention the license of the assets? [Yes] See Appx. E
  c. Did you include any new assets either in the supplemental material or as a URL? [Yes] Supplemental material.
  d. Did you discuss whether and how consent was obtained from people whose data you’re using/curating? [Yes] See Appx. E
  e. Did you discuss whether the data you are using/curating contains personally identifiable information or offensive content? [Yes] See Appx. E
5. If you used crowdsourcing or conducted research with human subjects…
  a. Did you include the full text of instructions given to participants and screenshots, if applicable? [N/A]
  b. Did you describe an y potential participant risks, with links to Institutional Review Board (IRB) approvals, if applicable? [N/A]
  c. Did you include the estimated hourly wage paid to participants and the total amount spent on participant compensation? [N/A]

## A Societal Impact Statement

Fitness estimation methods have wide potential for positive impact, including in understanding and diagnosing disease [19] and designing novel proteins that can accelerate research or treat disease [50]. However, like most powerful technologies, fitness estimation methods also have potential for negative impact. Improved ability to diagnose genetic disease could enable discrimination on the basis of genetic mutations, for instance insurance companies could refuse coverage or charge high premiums to patients with pathogenic variants. These risks can be mitigated via appropriate laws and regulations, such as the Genetic Information Nondiscrimination Act of 2008 in the US. Synthetic biology and molecular design are dual use technologies, and novel proteins could be designed that cause harm. For a further discussion of dual use concerns for machine learning-based molecular design, and recommendations for mitigation, see Urbina et al. [54].

## B Evolutionary dynamics models

Application of the Sella and Hirsh [49] model (Eqn. 1) in JFPMs rests on a number of assumptions; we briefly the most relevant here.

When applying Eqn. 1 to amino acid sequences, as is typical for fitness estimation models, we ignore biases that come from the genetic code, which can modify the steady state probability of amino acids (in the absence of fitness effects) away from a uniform distribution. This is justified practically by the small effect sizes: if at steady state an amino acid has probability 1*/*64 instead of 1*/*20, the total difference in log probability is log(1*/*20) − log(1*/*64) ≈ 1, which is small compared to (for instance) the log probability differences relevant for disease risk prediction with fitness models, which are ≈ 10 ([19], Extended data Fig. 3). Moreover, this bias only contributes an overall shift in amino acid probabilities, independent of position, and so does not change our main theoretical results. We ignore biases caused by asymmetric mutation rates for analogous reasons (though note they are often included in PMs in practice) [49].

The constant *β* depends on the effective population size, as well as the underlying population genetics model (Moran or Wright) and organismal ploidy ([49], Table 1). Following standard practice, we treat *β* as fixed for simplicity, though in reality it may vary over time and across lineages. Taking into account these possible changes clearly would not contradict our main theoretical result, that fitness and phylogeny are non-identifiable.

## C Proofs

### C.1 Proof of Proposition 2.3

*N*.*b. this result is known in the literature (see e*.*g. [22], Eqn. 1) but we are unaware of a proof, so we provide one here for completeness*.

*Proof*. For notational convenience, we will work with a standardized OUT, with *μ* = 0 and *σ* = 1. The final result can be obtained by translating and scaling the distribution of leaves. The transition distribution from point *x*′ at time *t*′ to point *X* at time *t* under the Ornstein-Uhlenbeck (OU) process is

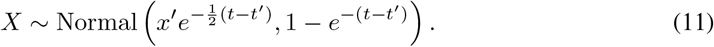

This distribution can be reparameterized in location-scale form as

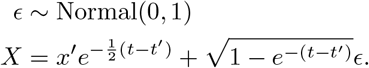

As *t* → ∞ we reach the stationary distribution Normal(0, 1). Let *b* ∈ {1, …, **B?** index the branches of the tree, let *λ*_*b*_ be the length of branch *b*, and let *j* ∈ {1, …, *N*} index the leaves (observed species or sequences); see Fig. 5. We have assumed that the most recent common ancestor of the observed sequences was sampled from *p*^∞^; this can be represented by adding a single branch length (indexed *b* = 1) to the root with length *λ*_1_ =. Let *ϵ*_*b*_ be the noise describing the OU diffusion over each branch. Let *ξ*_*j,b*_ be the total time from leaf *j* to the nearest vertex on branch *b*, so long as branch *b* is on the path from leaf *j* to the root; otherwise, set *ξ*_*j,b*_ = ∞. For instance, in the diagram in Figure 5, we have *ξ*_1,4_ = 0, *ξ*_1,2_ = *λ*_4_, *ξ*_1,1_ = *λ*_4_ + *λ*_2_, and *ξ*_1,5_ = *ξ*_1,6_ = *ξ*_1,7_ = *ξ*_1,3_ = ∞. We can now write the leaf position as

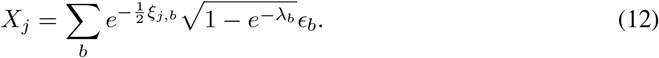

**Figure 5:**
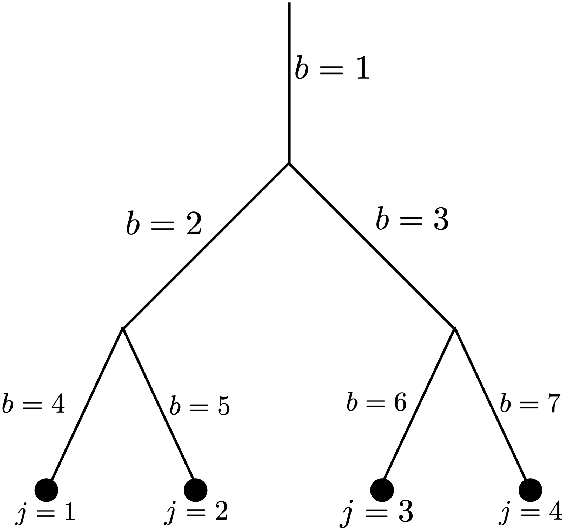
Tree labeling for the proof of Proposition 2.3

Define the matrix

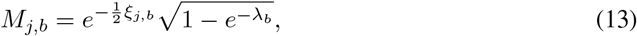

such that *X*_*j*_ = ∑ _*b*_ *M*_*j,b*_*ϵ*_*b*_. We can now describe the complete leaf distribution as

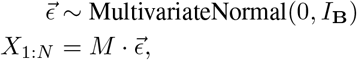

where *I*_**B**_ is the **B**-dimensional identity matrix. Thus, according to the location-scale representation of the multivariate normal,

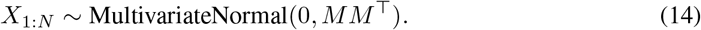

We can simplify the covariance matrix σ := *MM* ^⊤^. First

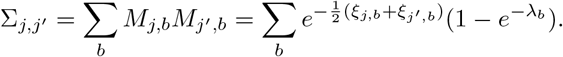

Before introducing the notation required to derive the general result, it’s helpful to get a sense of how the derivation works; in the example tree (Figure 5),

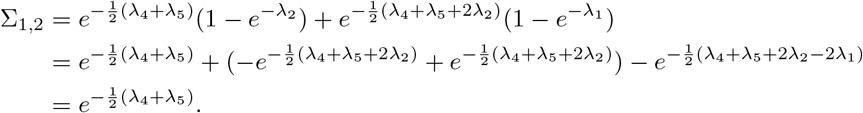

The sum over *b* telescopes, leaving only the initial term, which corresponds to the total time between leaf node 1 and leaf node 2. To construct the general result, define 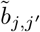 as the branch whose later node is the most recent common ancestor of leaves *j* and *j*′. In the example in Figure 6, 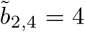.

**Figure 6:**
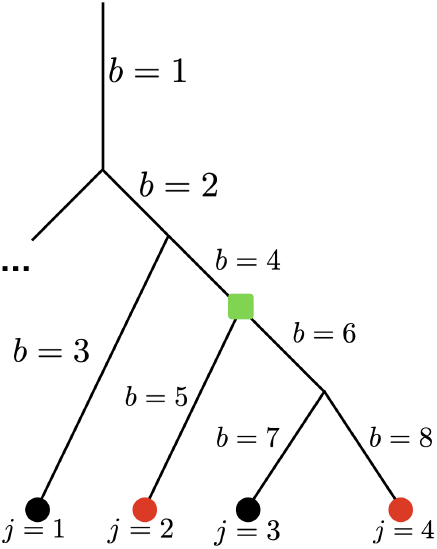
In red are the leaves considered in the examples in the proof of Proposition 2.3; in green is their most recent common ancestor.

**Figure 7:**
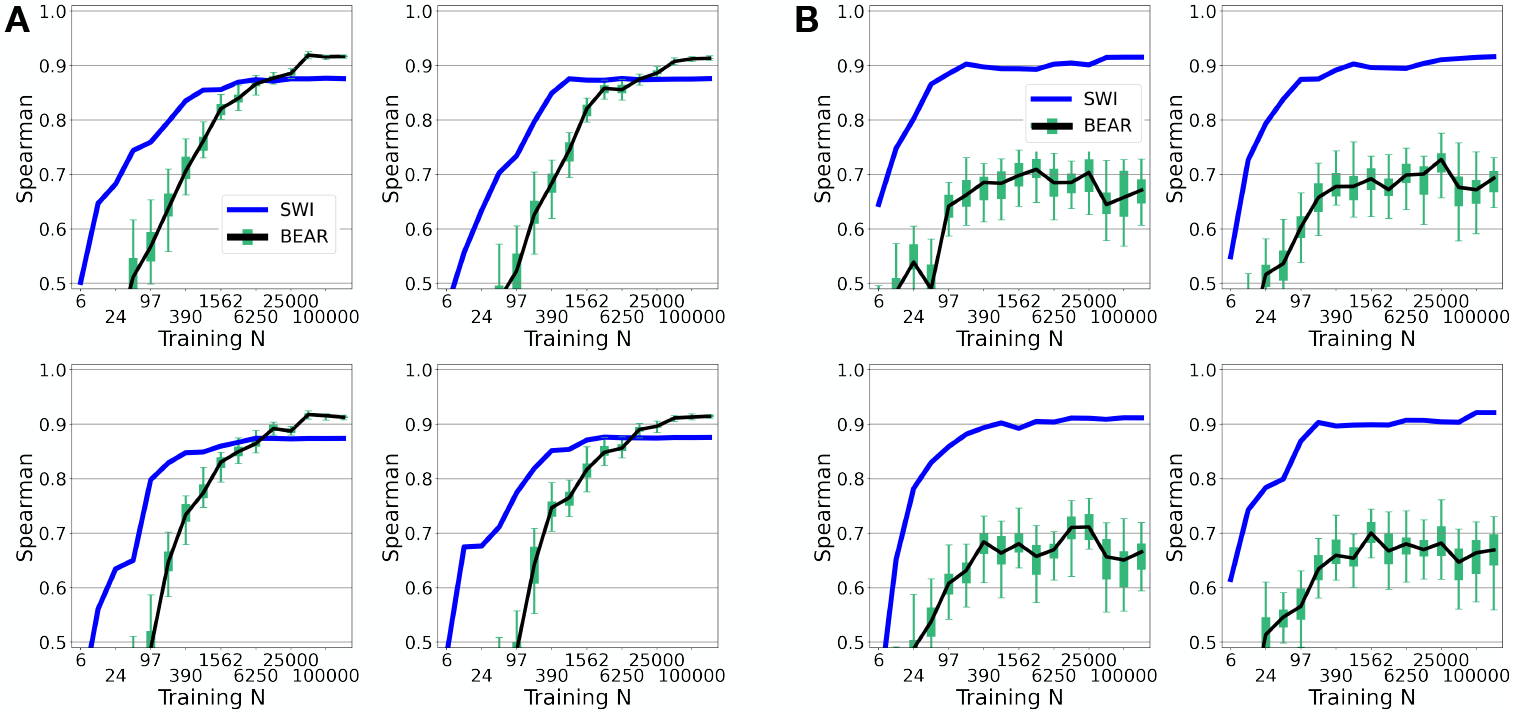
(A) Same as Fig. 3A, for four independent simulations following Scenario 1. (B) Same as Fig. 3B, for four independent simulations following Scenario 2.

**Figure 8:**
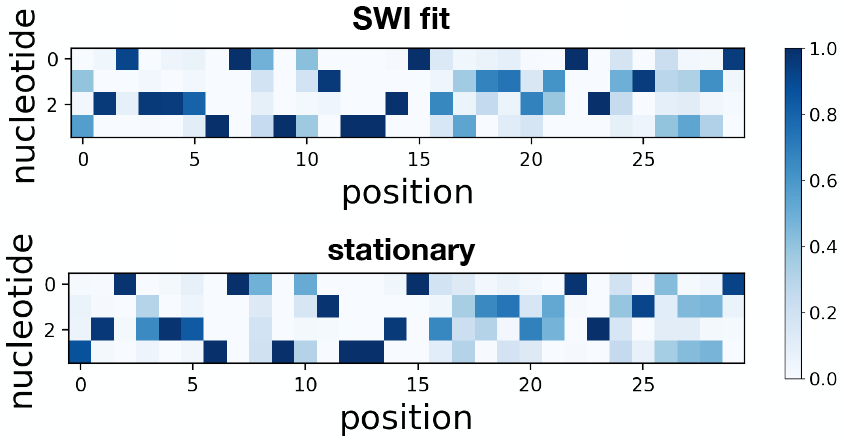
Probability of each nucleotide at each position learned by the SWI model (above) and in the stationary distribution *p*^∞^ (below), for a simulation from Scenario 2.

Let *R* be an ordered list of branches from 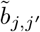 to *b* = 1, the earliest branch. In the example in Figure 6, *R* = [4, 2, 1]. Finally, let *t*_*jj*′_ be the length of the shortest path from leaf *j* to leaf *j*′, the time from the most recent common ancestor to *j* plus the time to *j*′. In the example in Figure 6, *t*_2,4_ = *λ*_5_ + *λ*_6_ + *λ*_8_. We now have

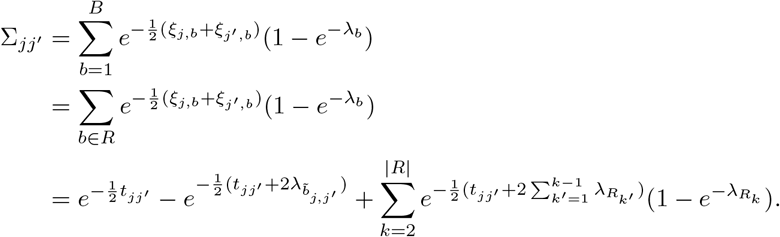

Breaking down the telescoping sum, and using the fact that the final element of *R* is *t*_1_ = ∞,

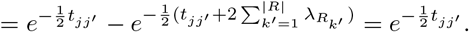

So we have the simple result that the covariance matrix depends just on the divergence times between leaves,

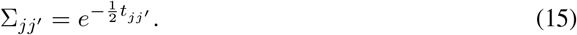

Translating the distribution Eqn. 14 by *μ* and scaling by *σ* yields the result.

### C.2 Proof of Theorem 3.3

Before proving the result, we briefly clarify a definition in the statement of the theorem:

#### Definition C.1

(Exchangeable in leaves). *Let* **H** *be a tree with countably infinite leaves and let* **H**_*π*_ *be a permutation of a phylogeny in its leaves, i*.*e. the same tree* **H** *with the leaves observed in a different order, according to a permutation π. A distribution over phylogenies is exchangeable in its leaves if p*(**H**) = *p*(**H**_*π*_) *for any permutation π*.

*Proof. Outline: First, using the results from Sarkar [48], we construct an embedding for each tree into the hyperbolic plane, being careful that the embedding preserves exchangeability. Second, we apply de Finetti*’*s Theorem to obtain the conditionally independent representation of the joint distribution of Z*_1_, *Z*_2_, *Third, we use the distortion bound from Sarkar [48] to bound the Wasserstein distance between p*(*ν*) *and* 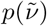.

First we describe the Sarkar [48] (1 + *ϵ*) distortion embedding algorithm setup. Vertices in phylogenetic trees have maximum degree three, and, by assumption, the minimum edge length in a tree **H** is greater than *η >* 0 with probability one. For any *ϵ*′ > 0, choose a *ρ < π/*3 and a scale factor

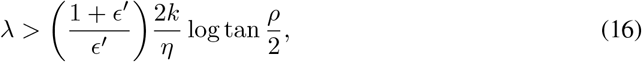

where *k* is the Gaussian curvature of the hyperbolic plane ℍ (for most hyperbolic geometry models, and in particular the Lorentz manifold, *k* = −1). Then, let *h*_1_(**H**), *h*_2_(**H**), … be the position of the leaves in the embedding of **H** produced by the (1 + *ϵ*) distortion embedding algorithm in Sarkar [48], using edge scale factor *λ*, and *ρ* separated cones with cone angle 2*π/*3 − 2*ρ*. Taking the last line of the proof of Theorem 6 in Sarkar [48], we are guaranteed that even for a countably infinite number of leaves,

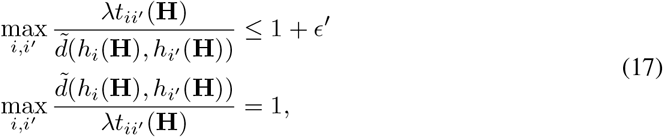

where *i, i*′ ∈ ℕ := *{*1, 2, …*}*, and 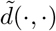 is the hyperbolic distance function.

Next we will modify the embedding function *h* to ensure that the distribution of embedded leaves is exchangeable. Let [**H**] be the set of phylogenetic trees that are equivalent to **H** up to reordering of the vertices. For each equivalence class [**H**] we choose one ordering of the vertices to be the canonical tree **Ĥ** ([**H**]), and for any tree **H** let *π*^*c*^(**H**) be the leaf permutation such that the reordered tree 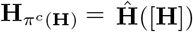. Now define the modified leaf embedding function 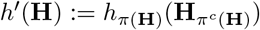 where *π*(**H**) is the inverse permutation of *π*^*c*^(**H**). Since by assumption the prior *p*(**H**) on the phylogenetic tree is exchangeable, we can rewrite *p*(**H**) using the induced distribution over equivalence classes *p*([**H**]) as

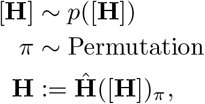

where Permutation is the uniform distribution over all permutations of ℕ := {1, 2,…}. We now define the distribution over leaf embeddings as

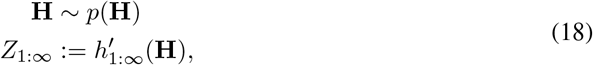

which we can rewrite as

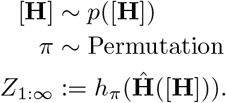

The distribution *p*(*Z*_1_, *Z*_2_, …) is therefore exchangeable. Applying de Finetti’s Theorem [29] we have a.s.

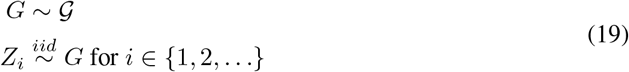

where *G* is a random measure distributed according to a prior 𝒢. Moreover, the embedding distortion bounds (Eqn. 17) are preserved for each **H**, since

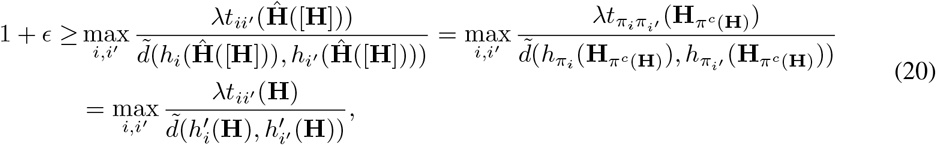

and by the same logic

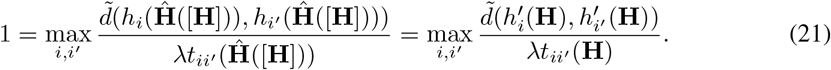

We will now construct the Wasserstein bound. Define the joint distribution over *ν* and 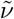,

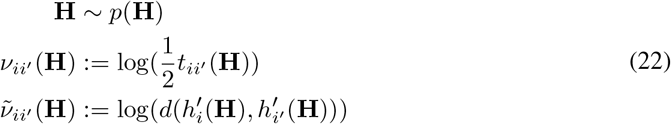

where we have chosen 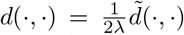. Note that the marginal distribution of *ν* matches its definition in the statement of the theorem, and that, applying Eqn. 18 and Eqn. 19, the marginal distribution of 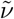 also matches its definition. Using the fact that log is a monotonically increasing function, Eqn. 20 gives

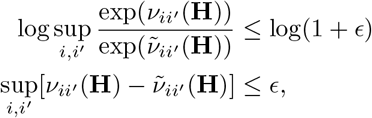

and similarly using the bound from Eqn. 21, 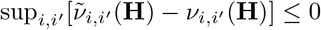. Thus, with probability 1 under *p*(**H**),

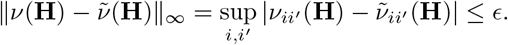

Recall that the Wasserstein distance between the distribution of two random variables *ν* and 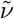 can be written as

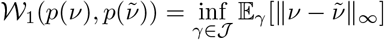

where 𝒥 is the set of joint distributions with marginals corresponding to the distributions of *ν* and 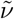 ([15], Chap. 11.8). Using the joint distribution in Eqn. 22, the Wasserstein distance is bounded by

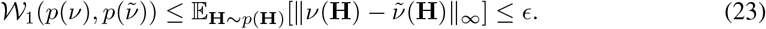

Now consider the case where 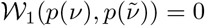. (N.b. in this case, we do not need to assume that the minimum time between nodes in **H** is greater than *η >* 0.) Since the Wasserstein metric is a metric on the space of probability distributions ([15] Lemma 11.8.3), 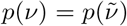 a.e.. Using the standard properties of Gaussian processes ([58], Chap. 2), the GPLVM model (Eqn. 5) can be written as

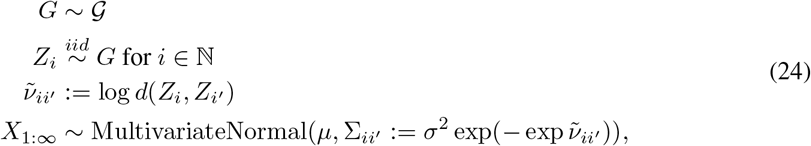

which is equivalent to the OUT distribution,

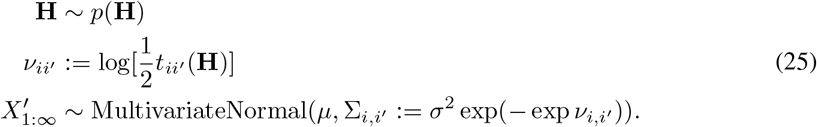

So the distribution *p*(*X*_1:∞_) produced by the GPLVM is equivalent to the distribution 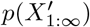 produced by the OUT model a.e..

### C.3 Proof of Theorem 4.1

We start with a more basic result that captures the intuition behind Thm. 4.1, and then prove a more general result, from which Thm. 4.1 can be derived as a corollary. In particular, we start by examining the special case where the model is well-specified with respect to *p*^∞^.

#### Proposition C.2.

*Assume that the model* ℳ *is log-convex and well-specified with respect to the stationary distribution, i*.*e. p*^∞^ ∈ ℳ. *Assume q*_*θ*_* *exists and is unique. Then, if the model is misspecified with respect to the data distribution, i*.*e. p*_0_ ∉ ℳ, *we have*

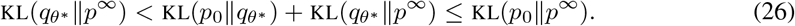

*But if the model is well-specified, i*.*e. p*_0_ ∈ ℳ, *we have*

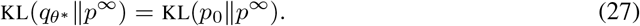

*Proof*. The first inequality in Eqn. 26 follows from the fact that ℳ is misspecified with respect to *p*_0_ and so KL(*p*_0_ ‖ *q*_*θ*_*) > 0. The second inequality follows from Thm. 1 from Csiszar and Matus [11], who study the geometry of reverse I-projections. For Eqn. 27, note that *q*_*θ*_* = *p*_0_ when *p*_0_ ∈ ℳ.

We next extend Prop. C.2 to the case where the model may be misspecified with respect to *p*^∞^ as well as *p*_0_.

#### Proposition C.3.

*Assume the model* ℳ *is log-convex and that q*_*θ*_* *exists and is unique. If*

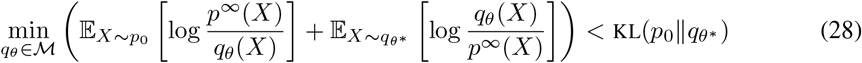

*then* KL(*q*_*θ*_* ‖ *p*^∞^) *<* KL(*p*_0_ ‖ *p*^∞^).

*Proof*. Define the projected stationary distribution

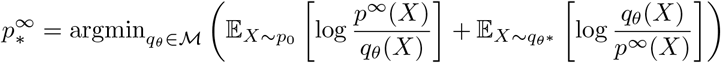

Now, from the definition of the KL divergence we have

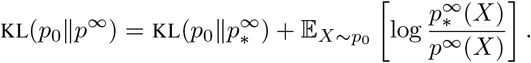

Applying Thm.1 from Csiszar and Matus [11] to KL 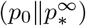 we have

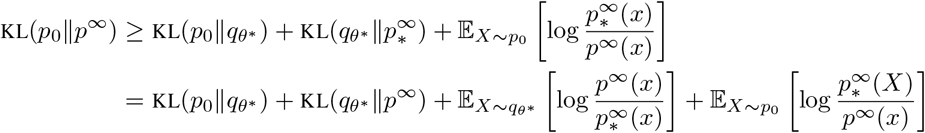

Applying Eqn. 28, the result follows.

One way of satisfying Prop. C.3 is for there to exist a 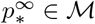 that is very close to *p*^∞^, in the sense that log 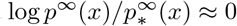 in areas of 𝒳 with high probability under both *p*_0_ and *q*_*θ*_*.

Finally we, derive the more interpretable (but also more restrictive) conditions in Thm. 4.1.

*Proof of* Thm. 4.1 For Eqn. 8, note that *q*_*θ*_* = *p*_0_ when *p*_0_ ∈ ℳ. To show Eqn. 7, we will show that the conditions of Prop. C.3 are met. (N.b. we will work with sums over *x* since we are concerned with discrete sequences, but the same derivation holds replacing sums with integrals.) We have

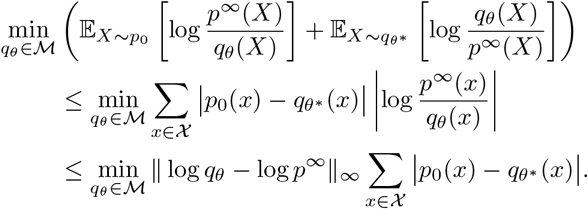

Applying Eqn. 6 to the first term and the definition of total variation distance to the second term,

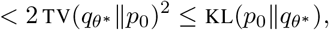

where the second inequality is Pinsker’s inequality. We see that Eqn. 28 is satisfied, and the result follows.

## D Simulation Details

In both scenarios, we generated sequences of fixed length |*X*| = 30, with an alphabet size of *B* + 1 = 4 (corresponding to nucleotides).

### Scenario 1

We simulated from a Potts model

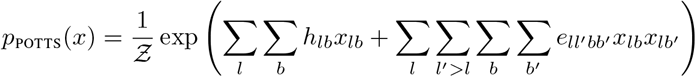

where *h* is the sitewise energies, *e* is the pairwise energies, *x* is a one-hot sequence encoding, *l* indexes sequence positions and *b* indexes letters. Following the simulations in Ingraham and Marks [27], which were intended to roughly match the statistics of typical real protein Potts models, we drew *h*_*lb*_ ∼ InvGamma(2, 0.8) and

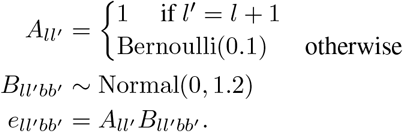

The energies *h* and *e* were drawn once, and the same values used across independent simulations. We sampled from the model using a Gibbs sampler with 100 steps of burn-in and 10 parallel chains using the code from Ingraham and Marks [27] (https://github.com/debbiemarkslab/persistent-vi). We shuffled the resulting samples to remove autocorrelation.

### Scenario 2

We used a site-wise independent fitness function:

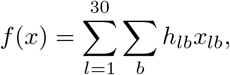

with site-wise residue biases *h*_*l*_, where *x*_*l*_ is a one-hot encoding of the letter at the *l*-th position of *x*. To generate phylogenetically correlated sequences, we sampled phylogenetic trees from a Kingman Coalescent ([3], Def. 2.1) with rate 1. Starting from a random sequence drawn from the steady state distribution at the root, we evolved the sequence simulating a Wright process in a haploid population ([49], Eqn. 3) according to the tree and fitness function. In particular, for sequences *x*_0_, *x* that are one mutation away, the mutation rate is

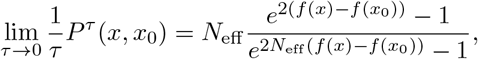

where we set the effective population size to *N*_eff_ = 10000. This stochastic process has steady state

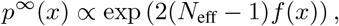

([49], Eqn. 7).

### SWI model

We fit the SWI model with maximum likelihood estimation.

### BEAR model

In these simulations, we used a vanilla BEAR model with a uniform embedded AR model (i.e. a Bayesian Markov model) for simplicity. We set the Dirichlet prior concentration to the constant *α* = 0.5. Based on the theoretical analysis in Amin, Weinstein and Marks [2] (Thm. 35), we used a prior on lags of the form

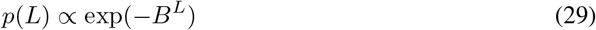

where *B* is the alphabet size (4 for nucleotides). We inferred the prior via empirical Bayes, marginalizing over the transition probabilities following the protocol in [2]. Conditional on lag *L*, sampling from the posterior over the BEAR model is straightforward thanks to Dirichlet-Categorical conjugancy.

### Evaluation

We defined 𝒮_*f*_ following standard protocols for fitness estimation models. In particular, we let 𝒮_*f*_ (*p*) be the Spearman correlation between *p*(*x*) and *f* (*x*) for *x* ∈ Λ where Λ consists of all possible single point mutations (i.e. single letter changes) of an initial (“wild-type”) sequence. The wild-type sequence was chosen as the most likely sequence under *p*^∞^, computed exactly for Scenario 2 and estimated based on the 10^6^ samples for Scenario 1.

To estimate model perplexity (Fig. 3C and 9B), we used *N* = 10, 000 independent sequences from *p*_0_ and computed the per-residue perplexity

**Figure 9:**
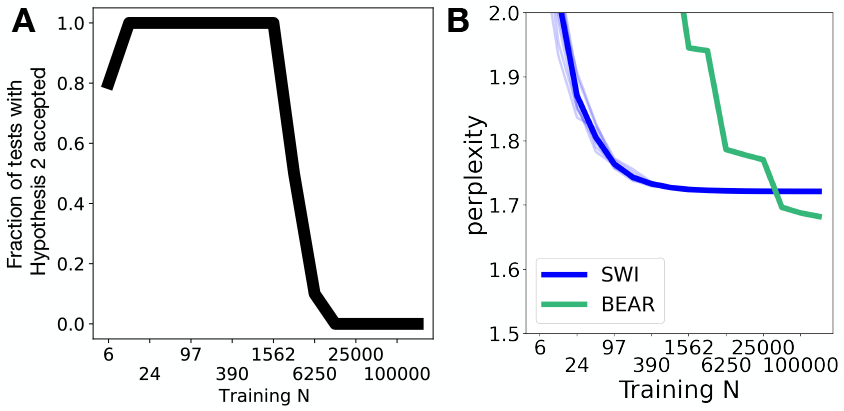
(A) Fraction of independent simulations (out of 10 total), following Scenario 1 (Sec. 6), in which Hypothesis 2 was accepted at level *α* = 0.025. (B) Perplexity on heldout data of the BEAR and the SWI models in Scenario 1. Thick line corresponds to the average over 10 individual simulations (thin lines).

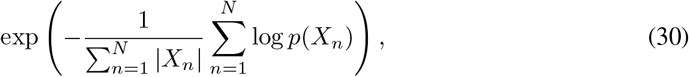

where |*X*_*n*_| is the sequence length and *p*(*X*_*n*_) is the probability of the sequence under the model.

To estimate the KL to the fitness distribution in Scenario 2 (Fig. 3D), we sampled *N* = 10, 000 independent sequences from *p*^∞^, {*X*_1_, …, *X*_*N*_ } and estimated

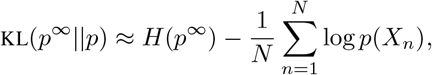

where *H*(*p*^∞^) is the entropy of *p*^∞^, which can be computed analytically. For BEAR, we plotted the KL to the posterior predictive, which, using Jensen’s inequality can also be seen to lower bound

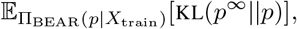

where Π_BEAR_(*p*|*X*_train_) is the BEAR posterior learned from the training dataset.

## E Empirical Results Details

### E.1 Data

#### Prediction task #1 (functional effect)

Following standard practice, we report the absolute value of the Spearman correlation as 𝒮_*f*_ (*p*), since in some assays a negative change in the measured quantity corresponds to larger fitness (note that in all cases the predicted directionality of the effect under each model was correct). We focused on single amino acid substitutions, taking only those for which EVE was able to make a prediction (EVE is limited by its reliance on a multiple sequence alignment). We used the same data as in Shin et al. [50], Table 1, taking the 37 experiments performed on the following 32 proteins: UBC9_HUMAN, UBE4B_MOUSE, P84126_THETH, HIS7_YEAST, BLAT_ECOLX, IF1_ECOLI, PTEN_HUMAN, B3VI55_LIPST, GAL4_YEAST, POLG_HCVJF, PABP_YEAST, CALM1_HUMAN, AMIE_PSEAE, TRPC_THEMA, RASH_HUMAN, YAP1_HUMAN, TRPC_SULSO, DLG4_RAT, BG_STRSQ, KKA2_KLEPN, HSP82_YEAST, B3VI55_LIPST (stabilized), MK01_HUMAN, HIV BF520 env, SUMO1_HUMAN, RL401_YEAST, PA_FLU, HG_FLU, TPMT_HUMAN, HIV BG505 env, TPK1_HUMAN, and MTH3_HAEAE (stabilized). The sequence data and assay data are publicly available for research use.

#### Prediction task #2 (pathogenicity)

We used the pathogenicity labels of single amino acid substitutions curated from ClinVar [31] in Frazer et al. [19]. Note ClinVar data is freely available for public use, and labels only depend indirectly on patient data; there is no per-person label, and no way of identifying individual patients [31]. We considered labels for 87 human proteins less than 250 amino acids in length: AICDA, AQP2, ATPF2, B9D2, CAH5A, CAV3, CD40L, CF410, CHC10, CIA30, CLD16, CLN8, COQ4, CRBB2, CRGD, CTRC, CXB1, CXB2, CXB3, CXB4, CXB6, CY24A, DERM, DGUOK, DHDDS, EDAD, EFTS, ELNE, ETFB, ETHE1, EXOS3, FGF10, FGF23, FOXE3, FRDA, GP1BB, HBB, HEM4, HSPB1, HSPB8, IFM5, IFT27, JAGN1, KAD2, KCNE1, KCNE2, KITM, LITAF, MMAB, MMAC, MPU1, MYPR, NDP, NDUS8, NFU1, NKX25, NMNA1, OPA3, PAHX, PDYN, PMM2, PMP22, PNPH, PNPO, PROP1, PSPC, PTPS, RASH, RNH2A, S5A2, SAP3, SBDS, SCO1, SDHB, SDHF2, SIX1, SIX3, SOMA, TMM70, TNNT2, TPK1, TPM2, TR13B, TWST1, VHL, XLRS1, ZC4H2.

#### Training data

All models were trained on datasets of protein sequences gathered as described in [50] for pathogenicity effect prediction tasks and as described in [19] for functional effect prediction tasks. SWI and EVE were trained on the multiple sequence alignment, while Wavenet and BEAR were trained on the raw sequences as described in [50]. All datasets were uniformly subsampled to produce a 75%/25% train/test split.

#### E.2 Models and code

The SWI model was trained via maximum likelihood.

The Wavenet model was trained via maximum likelihood with the default architecture, hyperparameters and training protocol described in [50], for 100,000 steps. Code is from https://github.com/debbiemarkslab/SeqDesign published under the MIT licence. We did not apply the Wavenet model to the second prediction task, as it has only previously been developed for the first task.

The EVE model was trained via variational inference, using the same architecture, hyperparameters, and training protocol described in [19]. Code is from https://github.com/debbiemarkslab/ EVE published under the MIT licence. To match the protocol of the original paper, EVE was – unlike SWI, Wavenet and BEAR – (a) trained on the full dataset rather than the training set alone, and (b) used a sequence reweighting heuristic.

The BEAR model used an embedded convolutional neural network (the same architecture as used in [2], with layer 1 width of 16, filter width of 5 and 30 filters total) and a uniform prior over lags 2, 3, 5, 7, and 9. Code is from https://github.com/debbiemarkslab/BEAR published under the MIT licence. The model was trained using empirical Bayes, as described in [2], for 500 steps with a batch size of 500000 kmers. Two Nvidia TeslaV100s GPUs were used, on an internal cluster; training took up to 8 hours. To construct posterior credible intervals, we used 41 samples from the posterior for prediction task #1, and 1000 samples for prediction task #2.

To train and predict using BEAR one needs to transform sequences into de-Bruijn graphs. To do so for amino acid sequences, we used code from https://github.com/jdisset/kmap. We communicated with the author of this code to obtain permission for its use.

We computed the heldout perplexity (Eqn. 30) for the BEAR posterior predictive and for Wavenet to produce Fig. 10.

**Figure 10:**
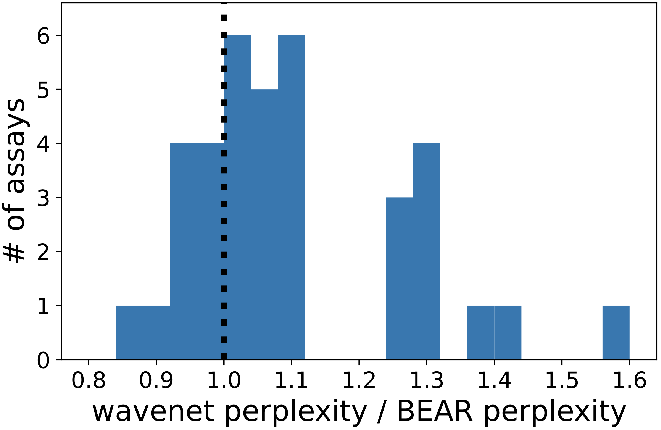
Ratio of the per residue perplexity on heldout data of the Wavenet model and of the BEAR model posterior predictive, across the 37 assays used for the first prediction task. Note lower perplexity corresponds to better density estimation performance.

### E.3 Convergence experiments

To plot the convergence of the posterior over *p*_0_ as a function of *N* (Fig. 4CD, 11 and 12), we used a vanilla BEAR model, a nonparametric Bayesian Markov model. Note that here we fixed the embedded AR model, rather than refitting with larger *N*, so that we could analyze the the convergence behavior with reference to the asymptotic results of Thm. 35 in [2], which does not take into account empirical Bayes. We set the Dirichlet concentration to 10 and used a prior over lags as in Eqn. 29.

**Figure 11:**
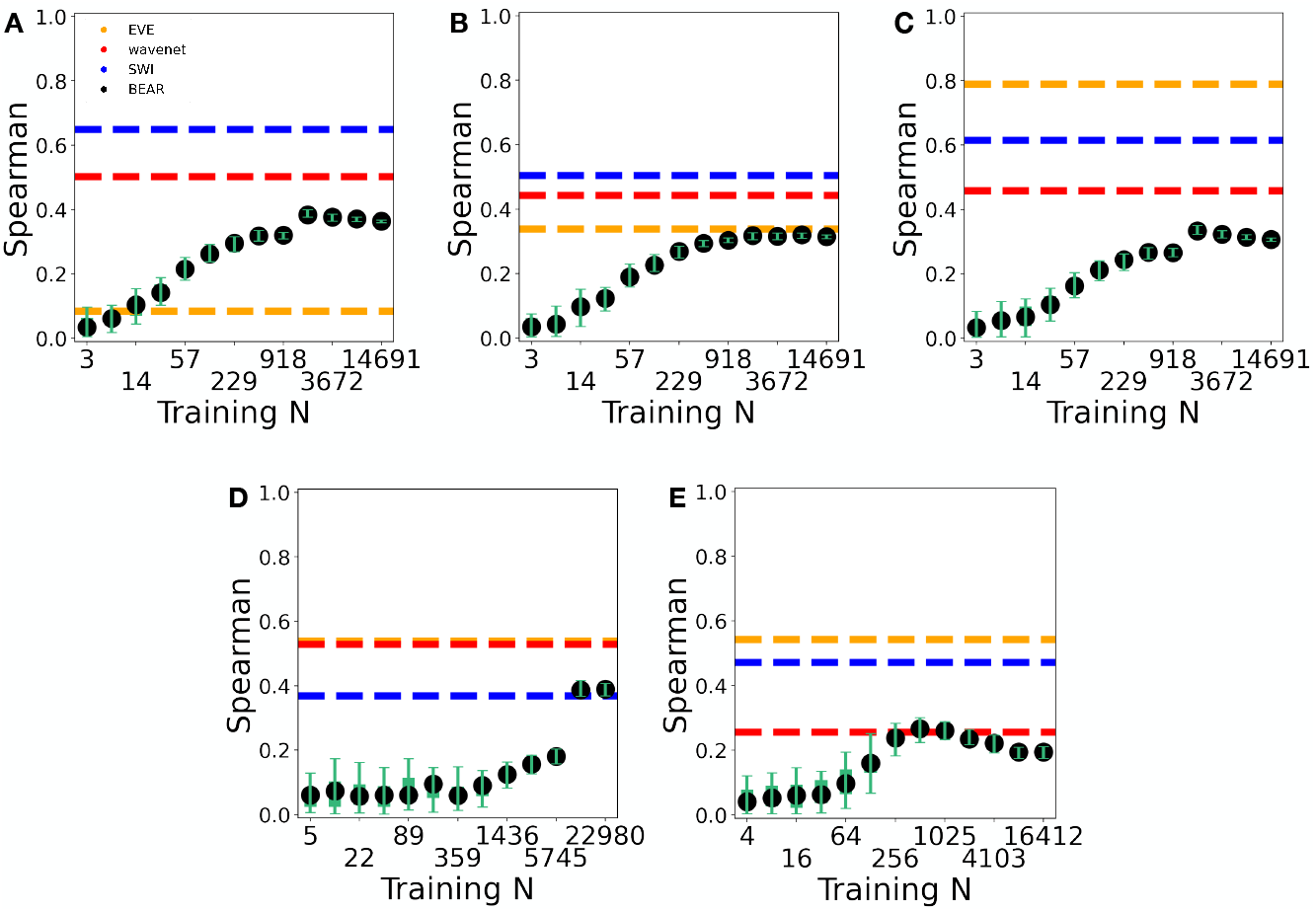
Same as Fig. 4CD, for 5 additional assay examples. A-C are each distinct *β*-lactamase assays; D is from GAL4 (DNA-binding domain); E is from UBE4B (U-box domain).

**Figure 12:**
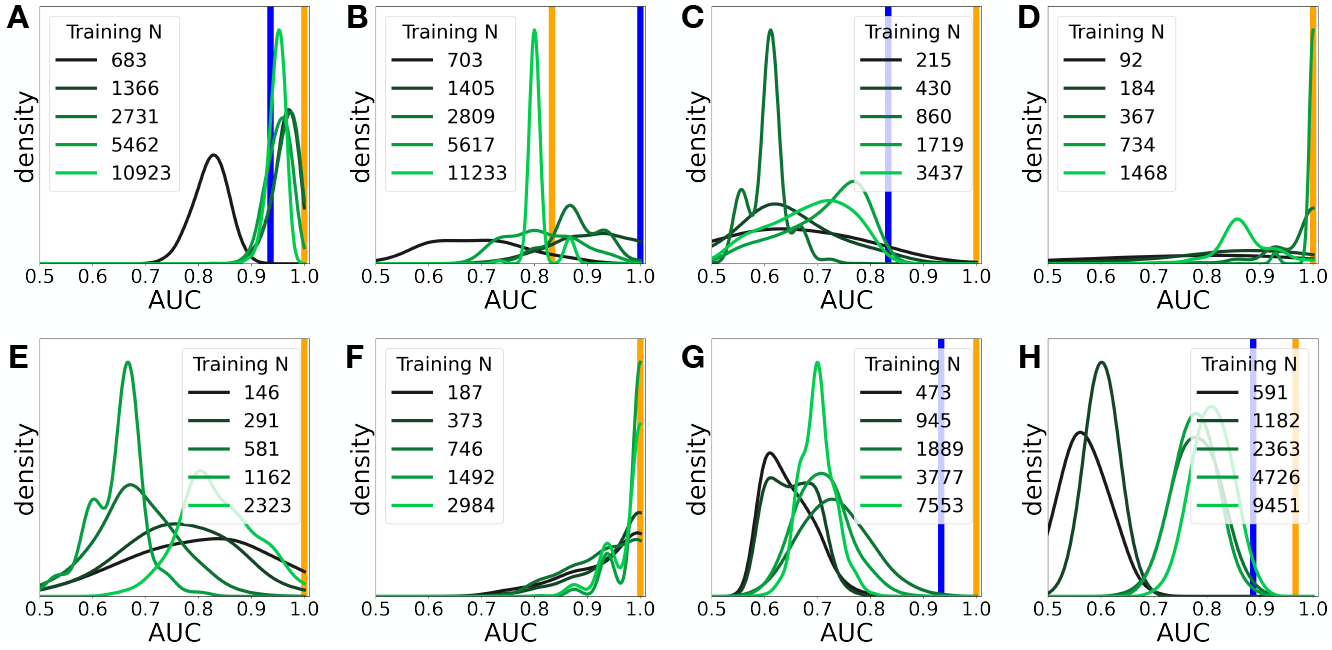
Convergence of the BEAR posterior over AUCs with *N* (green distributions), compared to the AUC of SWI (blue line) and EVE (yellow line), for the second prediction task. (A) is for the *CXB1* gene, (B) *CXB6*, (C) *EXOS3*, (D) *FGF23*, (E) *OPA3*, (F) *PAHX*, (G) *PROP1*, (H) *S5A2*.

### E.4 Interpolation experiments

We fit a BEAR model using the architecture and training protocol described in Sec. E.2, optimizing both the parameters of the AR model and *h* via empirical Bayes. We then varied *h* from its optimized value, and recalculated the total marginal likelihood and the posterior distribution over 𝒮_*f*_ (*p*) (Fig. 4EF and 13). We also computed the value of 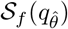 for the fit BEAR model in the *h* → 0 limit (Fig. 14).

**Figure 13:**
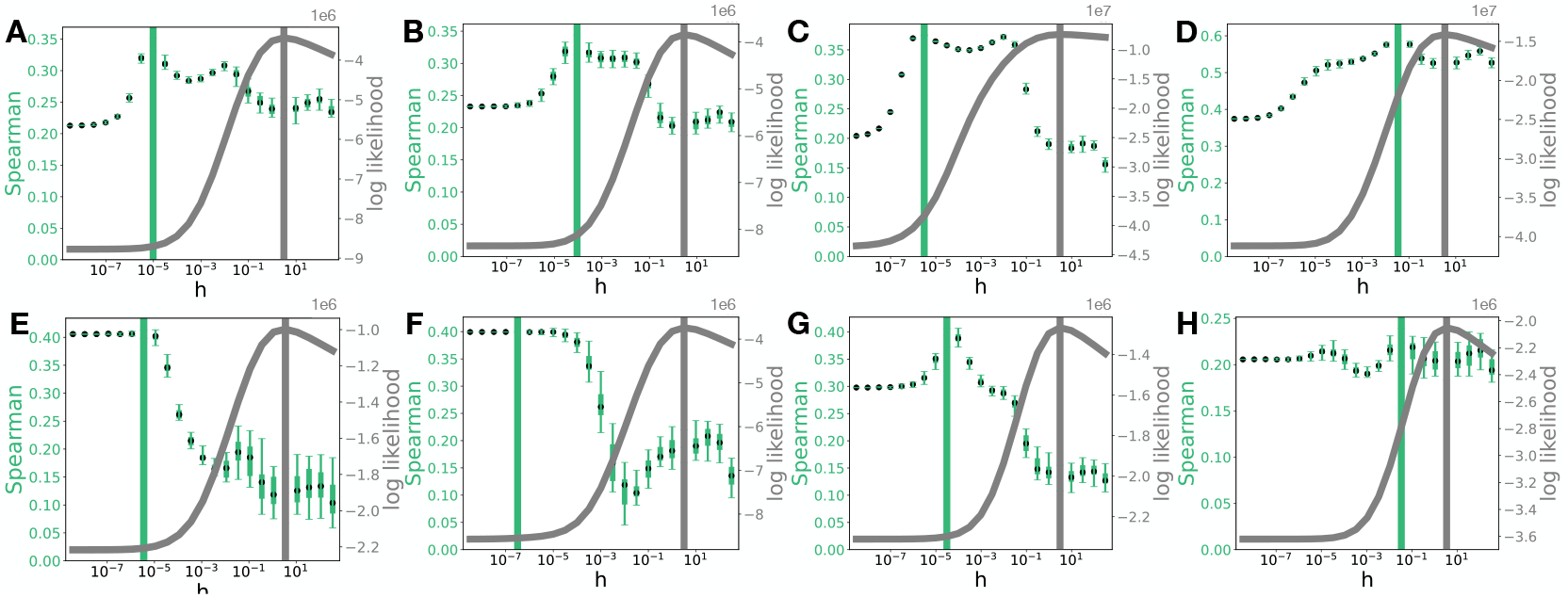
Same as Fig. 4EF, for 8 additional assay examples. (A) Aliphatic amidase, (B) levoglucosan kinase (stabilized), (C) HIV env protein (BF520), (D) *β*-glucosidase, (E) UBE4B (U-box domain) (F) TIM barrel, (G) thiopurine S-methyltransferase, (H) thiamin pyrophosphokinase 1.

**Figure 14:**
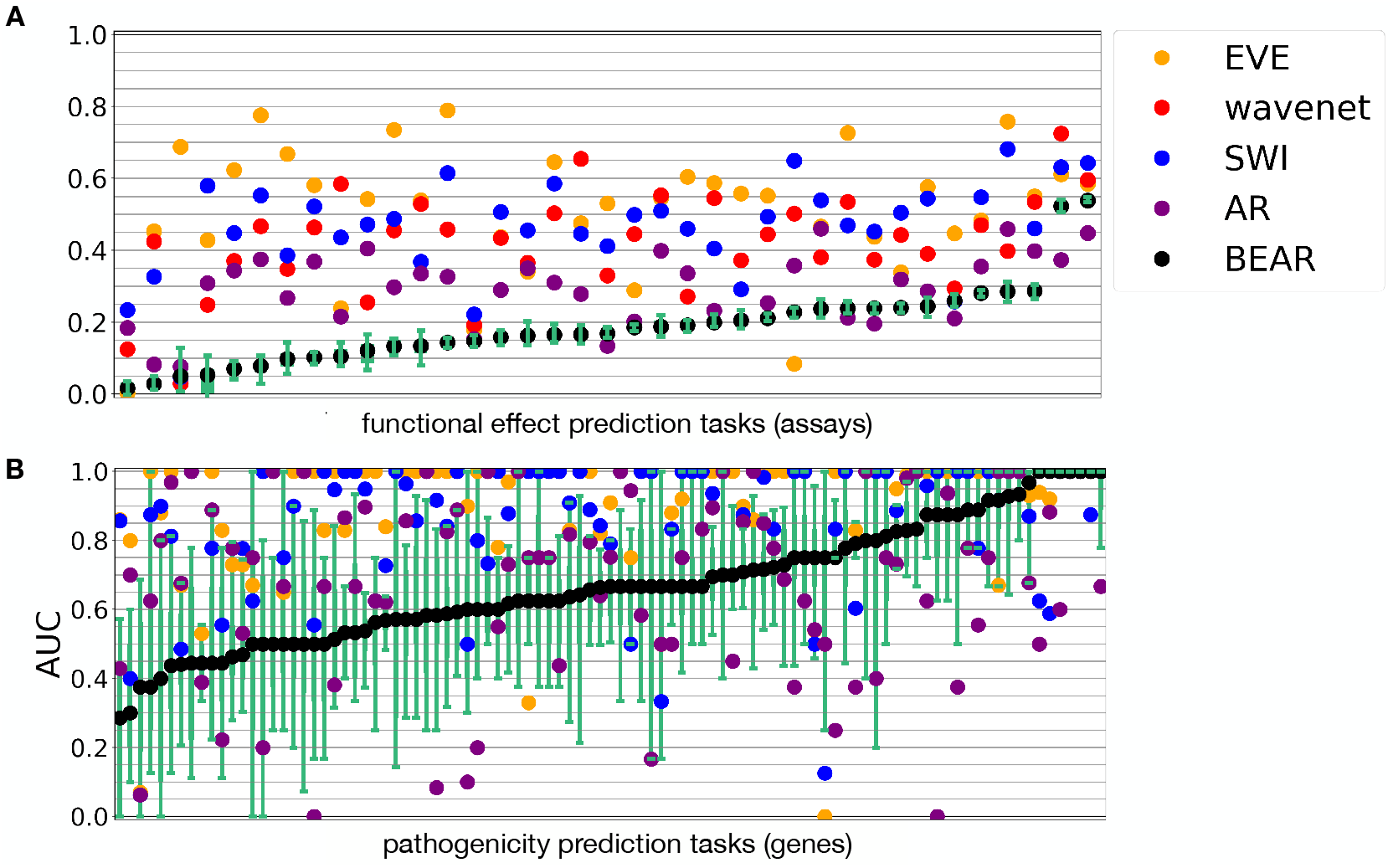
Same as Fig. 4AB, with the addition of the AR model in the *h* → 0 limit (purple). In prediction task 1 (A), Hypothesis 2 is accepted in 28/37 assays (75%) while Hypothesis 1 is accepted in 6/37 (16%) for the AR model. In prediction task 2 (B), Hypothesis 2 is accepted in 16/97 genes (16%) and Hypothesis 1 is accepted in 17/97 genes (18%) for the AR model.

## F Supplementary code

The supplementary code provides a Jupyter notebook (example.ipynb) illustrating the application of our BEAR diagnostic test on simulated data. It is available under an MIT License.

## G Discussion of Protein Language Models

In this section we discuss the relevance and relationship of our results to large-scale “protein language models” such as ESM-1v [36], MSA Transformer [43], UniRep [1], Tranception [41], ProGen [33], ProGen2 [40] and others. Note the term “protein language model” is something of a misnomer; these methods are far from unique in applying and extending ideas from natural language processing (NLP) to build generative protein sequence models (Wavenet [50] and BEAR [2] being just two other examples). Rather, inspired by the recent success of large-scale “foundation models” in NLP, protein language models have two key distinguishing properties: (1) they use neural network architectures with very large numbers of parameters and (2) they are trained on very large and very diverse datasets, not just sequences from a single family [4].

Our results suggest that better density estimation for sequence distributions does not necessarily imply better fitness estimation, and that a carefully chosen misspecified model can systematically outperform well-specified models. We believe our results complicate but *do not contradict* lessons learned on the importance of scale in natural and protein language models.

### Big and Diverse Data

Protein language models are trained on datasets that are bigger than those used for single family models, not only in the sense of containing more sequences but also in the sense of being more diverse. In other words, they differ not only in *N* but in the data generating distribution *p*_0_. The idea that using such big and diverse datasets may lead to better fitness estimation is entirely in line with our theory. First, *given* a particular model class ℳ and *p*_0_, we expect datasets with larger *N* to be better, as it brings our estimated distribution 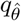 away from the prior and closer to the projected distribution *q*_*θ*_*. Second, while increasing the diversity of *p*_0_ does not remove the non-identifiability problem, it can enable practical strategies for reducing the effects of phylogeny. Misspecified models fit to many different protein families may be more robust to the “noise” introduced by phylogeny, as they must be accurate across families with very different phylogenetic structure. Moreover, protein language models are often trained on datasets that have been thinned, with very similar sequences removed (e.g. UniRef90 [36] rather than UniRef100 [44]); this shifts the target distribution *p*_0_, and can potentially ameliorate the effects of phylogeny.

### Downstream Tasks

The problem of fitness estimation using generative sequence models can be thought of as solving a downstream task of predicting an external label by learning an unsupervised one-dimensional representation for each sequence (namely, its log likelihood under the model).^3^ In general, for problems that involve using unsupervised representations for downstream tasks, progress often comes from directly optimizing the hyperparameters of the unsupervised model to improve downstream task performance, without regard to whether such hyperparameter changes increase or decrease density estimation performance of the unsupervised model. Our results suggest that focusing on optimizing hyperparameters and architectures for downstream performance instead of density estimation, perhaps by using meta-learning, may also be a successful way forward for fitness estimation [37].

### Architecture Choices

Our results are most directly in tension with recommendations from NLP for model size: we show theoretically and empirically that in the large *N* limit, misspecified models can be more accurate fitness estimators than well-specified models, which can conflict with the standard recommendation in large-scale natural and protein language models to give models as many parameters as possible. Our prediction is born out in the recent results of Nijkamp et al. [40], which demonstrate that a protein language model with more parameters and better density estimation can be worse at fitness estimation.

Still, a systematic understanding of model size and its tradeoffs in protein language models is lacking. For example, it may be advantageous to use misspecified models which have a very large numbers of parameters, as high-parameter models can possess other desirable properties beyond expressiveness, such as smoothness [7]. Indeed, many modern neural architectures (including attention layers in transformers) enforce invariance to certain symmetries (such as permutation or translation), ensuring that they are not universal approximators even when they have very large numbers of parameters [5]. Regularization and early stopping can also effectively restrict the model class ℳ.^4^ Broadly, our results fit with the idea that neural architecture and hyperparameter choice can substantially impact downstream tasks even in an age of massive models and data.

The question of whether protein language models can systematically outperform the true data distribution *p*_0_ at fitness estimation remains an open problem for future work. Here we describe some of the difficulties in addressing the problem and present preliminary results. The first challenge is scaling the diagnostic test to massive datasets. It is crucial, in particular, to work with datasets with large enough *N* that the test has power to reject the null hypothesis; convergence of the BEAR diagnostic is expected to be slower with (roughly speaking) high diversity *p*_0_ (Thm. 35, [2]). Computationally, scaling the test to massive datasets will require the use of large scale amino acid kmer counters, which are currently less performant than nucleotide kmer counters [30]. A second challenge is evaluating the density estimation performance of protein language models. Models trained using masked language modeling objectives (such as ESM-1v [36] and MSA Transformer [43]) do not specify a probability distribution over full sequences, i.e. they do not estimate a well-defined density [20]. Moreover, in some cases ultra-large scale models and/or training procedures are kept private or semi-private, making it difficult to understand what exactly was in the training and test sets [33, 6].

Nevertheless, we can rule out the idea that protein language models all achieve excellent fitness estimation and excellent density estimation on single family datasets. We studied UniRep, an LSTM-based protein language model trained on a diverse evolutionary dataset (UniRef50), since UniRep provides a well-defined density over sequences (unlike ESM-1v and MSA Transformer) and is publicly available (unlike ProGen) [1]. We evaluated UniRep on the experimental assay prediction task, comparing to the BEAR posterior predictive distribution and to Wavenet, both of which were inferred from single family datasets (as described in Sec. 7). If UniRep were to show similar or better perplexity to the BEAR posterior predictive, and similar or better fitness estimation performance to Wavenet, it could potentially represent a counterexample to our theory on the benefits of misspecification. Instead, we find that UniRep has substantially worse perplexity as compared to the BEAR posterior predictive (Fig. 15) and similar fitness estimation performance to the BEAR posterior predictive (median Spearman of 0.174 for UniRep, 0.169 for BEAR and 0.443 for Wavenet). We therefore see no evidence that protein language models contradict our findings on single family models, though much further work remains to be done.

**Figure 15:**
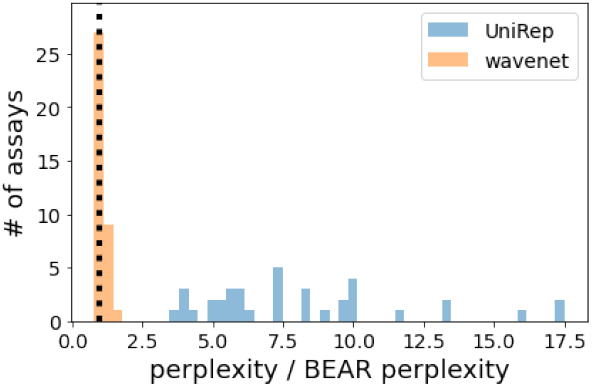
Ratio of the per residue perplexity on heldout data of the UniRep and Wavenet model and of the BEAR model posterior predictive, across the 37 assays used for the first prediction task. Note lower perplexity corresponds to better density estimation performance.

## Footnotes

3 N.b. since the representation is one-dimensional, and the downstream predictor is assumed to be a monotonically increasing function, success can be evaluated in a “zero-shot” setting using Spearman correlation or AUC, instead of training a supervised predictor on a small amount of labelled data.

4 In particular, this can occur if the amount of regularization or the point of early stopping change with increasing amounts of data *N*, say because these values are chosen based on performance on a downstream task. In this case, regularization and early stopping will not act like a standard parametric prior, and their effects will not necessarily be asymptotically washed out in the large *N* limit.

