## Supplemental code for "Non-identifiability and the Blessings of Misspecification in Models of Molecular Fitness": bear_preprocess.py

```
import numpy as np

from bear_model import dataloader
from bear_model import core

import random
import os
import tensorflow as tf

def dump_seqs_func(seqs, fname, alphabet_name):
    """Dump one-hot encoded sequences into fasta files"""
    alphabet = core.alphabets_en[alphabet_name][:-1]
    seqs_alph = [''.join(seq) for seq in 
                 alphabet[np.argwhere(seqs)[:, -1].reshape(np.shape(seqs)[:2])]]
    with open(fname, 'w+') as f:
        for i, seq in enumerate(seqs_alph):
            f.write('>{}\n{}\n'.format(i, seq))

def get_kmers(fname, lag, max_lag, alphabet, out_folder='seqs/'):
    """Preprocess the above sequences if not already done so and then get their kmer counts as a tensorflow data object.
    Uses summarize preproccesing function from BEAR."""
    rand_num = random.randint(0, 1e9)
    temp_folder = os.path.join(out_folder, 'tempf_{}'.format(rand_num))
    temp_file_loc = os.path.join(out_folder, 'temp_{}.txt'.format(rand_num))
    out_file_pref = os.path.join(out_folder, '{}_kmers'.format(fname))
    out_file_name = '{}_kmers_lag_{}_file_0.tsv'.format(fname, lag)
    if out_file_name not in os.listdir(out_folder):
        # Create a temporary file that describes the format for preprocessing
        with open(temp_file_loc, 'w+') as f:
            f.write('{},0,fa\n'.format(os.path.join(out_folder, 'train_{}.fa'.format(fname))))
            f.write('{},1,fa\n'.format(os.path.join(out_folder, 'test_{}.fa'.format(fname))))
            f.write('{},2,fa'.format(os.path.join(out_folder, 'infty_{}.fa'.format(fname))))
        os.system('mkdir ' + temp_folder)
        os.system('summarize.py {} {} -l {} -mk 18 -mf 1000 -t {}'.format(
            temp_file_loc, out_file_pref, max_lag, temp_folder))
        os.system('rm -r ' + temp_folder)
        os.system('rm ' + temp_file_loc)
    return dataloader.dataloader(os.path.join(out_folder, out_file_name), alphabet, 500000, 3)
            
def preprocess(train_seqs, test_seqs, true_seqs, fname, lag, max_lag,
               alphabet='dna', bear_prior=[0.5], dump_seqs=True, dump_seqs_folder='seqs'):
    """Perform entire pipeline of dumping sequence files, preprocessing them and calculating likelihoods"""
    # Dump sequences to preprocess into separate fasta files
    if dump_seqs:
        dump_seqs_func(train_seqs, os.path.join(dump_seqs_folder, "train_{}.fa".format(fname)), alphabet)
        dump_seqs_func(test_seqs, os.path.join(dump_seqs_folder, "test_{}.fa".format(fname)), alphabet)
        dump_seqs_func(true_seqs, os.path.join(dump_seqs_folder, "infty_{}.fa".format(fname)), alphabet)

    data = get_kmers(fname, lag, max_lag, alphabet, out_folder=dump_seqs_folder)
    
    # Calculate the likelihoods of the data under vanilla BEAR from the kmer counts
    pairs = tf.convert_to_tensor([[1, 1, 0], [1, 0, 1], [1, 0, 0]], dtype=tf.float64)
    data_lik = data.map(lambda a, b: tf.einsum('lj, ijk -> ilk', pairs, b))    
    (test_and_train_log_lik, infty_and_train_log_lik, train_log_lik) = dataloader.bmm_likelihood(data_lik, np.array(bear_prior))
    # Condition likelihoods
    test_log_lik_under_train = test_and_train_log_lik - train_log_lik
    infty_log_lik_under_train = infty_and_train_log_lik - train_log_lik
    return  (test_log_lik_under_train, infty_log_lik_under_train, train_log_lik)
```
