## Supplemental code for "Non-identifiability and the Blessings of Misspecification in Models of Molecular Fitness": example.ipynb

```
# %% [markdown]
This notebook is associated with the paper "Non-identifiability and the Blessings of Misspecification in Models of Molecular Fitness and Phylogeny". It recreates one replicate of the results in Figures 4 B, 4 C, 4 D, and S4 in the paper, demonstrating the blessings of misspecification when learning from phylogenetically correlated data.

# %% [markdown]
Dependencies:
1) The accompanying KingmanCoalescent.py file (code for sampling from a JFPM with a Kingman coalescent prior and site-wise independent fitness function).
2) The accompanying bear_preprocess.py file (code to calculate likelihoods of sequence datasets under the vanilla BEAR model).

Installation:
1) Use a python 3.8 environment with jupyterlab.
2) Install the ete3 python module, e.g. with ``pip install ete3``
3) Install the matplotlib python module, e.g. with ``pip install matplotlib``
4) Install the BEAR python module. The repository is at https://github.com/debbiemarkslab/BEAR, installation instructions are at https://bear-model.readthedocs.io/en/latest/installation.html. 

# %% [markdown]
First we set up the parameters of our simulation

# %% [code]
import numpy as np
from scipy.special import logsumexp
from bear_model import core

np.random.seed(1)
# Load the DNA alphabet
alphabet_name = 'dna'
alphabet = core.alphabets_en[alphabet_name][:-1]

# We'll be generating n_train sequences to train on
n_train = 100000
# We'll use n_test sequences to calculate perplexities
n_test = 10000
# We'll use n_true sequences from p^\infty to calculate KLs
n_infty = 10000
# For this tutorial, we'll run one replicate
replicates = 1
# Get n_cuts datapoints for number of training points (each half the amount of data as the next)
n_cuts = 16

# set up parameters for BEAR
# get mc_samples from bear
mc_samples = 41
#set the parameter in the lag prior of vanilla BEAR
c = 1
# Set the transition prior parameter of vanilla BEAR
bear_prior = 0.5
# We will train BEAR up to max_lag. The MAP lag will likely not be over 6 when c=1.
# In the interest of time we will assume this is the case in this tutorial; we trained models up to lag 13 in the manuscript. 
max_lag = 6
lags = np.arange(max_lag)+1

# Set the prior parameter on the SWI models we will train
swi_prior = 0.5

# Parameters for pylogenetic sequence generation
# Effective population size
N = 10000
# Length of the sequence
L = 30
# length of the alphabet
B = len(alphabet)
# Mutation rate
mu = 1

# %% [markdown]
Now we generate and save the sequences from $p_0$ (this may take a few minutes)

# %% [code]
from KingmanCoalescent import sample_coalescent, sample_pSH

# First we generate our phylogenetically corellated sequences
# First we sample a tree
coalescent = sample_coalescent(n_train*replicates+n_test)
# Then we choose p^\infty (h) and generate sequences along the tree according to the fitness.
sequences, h = sample_pSH(coalescent, L, B, pop_size=N, rate=mu)

# We shuffle our sequences and train/test split
np.random.shuffle(sequences)
train_seqs = [sequences[l:l+n_train] for l in np.arange(0, n_train*replicates, n_train)]
test_seqs = sequences[-n_test:]

# %% [markdown]
Next we calculate and sample from $p^\infty$

# %% [code]
# Calculate p^\infty (see supplementary section C)
nu = 2 * (N - 1)
steady_state = np.exp(-nu * h - logsumexp(-nu * h, axis=-1, keepdims=True))
log_ss = np.log(steady_state)
infty_ent = np.einsum('lb,lb->', steady_state, np.log(steady_state))

# generate sequences from p^\infty
infty_sequences = []
for l in range(L):
    infty_sequences.append(np.random.multinomial(1, steady_state[l], size=n_infty)[:, None, :])
infty_sequences = np.concatenate(infty_sequences, axis=1)

# %% [markdown]
Now we calculate mutation probabilities and define a function to give us spearmans from mutation probabilities.

# %% [code]
# Define the wt sequence as the most likely under p^\infty
wt_seq_inds = np.argmax(steady_state, axis=1)
# in letters
wt_seq = ''.join(alphabet[wt_seq_inds])
# one-hot encoded
wt_seq_ohe = wt_seq_inds[:, None] == np.arange(B)
# Make an array of all mutations in the format A10C for example.
vars_ = ['{}{}{}'.format(alphabet[wt_aa], i, alphabet[mt_aa]) for i, wt_aa in enumerate(wt_seq_inds) for mt_aa in np.arange(B) if mt_aa != wt_aa]

# get true mutation probabilities
def get_mut_probs(mut_mat):
    """From a matrix of log probabilities of mutant sequences, get an array of mutation log probabilities"""
    return (mut_mat - mut_mat[np.arange(L), wt_seq_inds][:, None])[~wt_seq_ohe]
true_mut_probs = get_mut_probs(log_ss)
num_muts = len(true_mut_probs)

# %% [markdown]
Now we will preprocess and analyze subsets of the training data. This will take a few minutes.

# %% [code]
from bear_preprocess import preprocess, get_kmers
from bear_model import get_var_probs

%load_ext autoreload
%autoreload 2

# a folder to output BEAR models
! mkdir seqs

probs_swi_test = np.empty([replicates, n_cuts])
probs_swi_infty = np.empty([replicates, n_cuts])
probs_swi_muts = np.empty([replicates, n_cuts, num_muts])

probs_bear_test = np.empty([replicates, n_cuts, max_lag])
probs_bear_infty = np.empty([replicates, n_cuts, max_lag])
probs_bear_train = np.empty([replicates, n_cuts, max_lag])
probs_bear_muts = np.empty([replicates, n_cuts, max_lag, num_muts, mc_samples])

# Loop through replicates
for r, train_seqs_set in enumerate(train_seqs):
    # loop through how much to cut the training data
    for i in range(n_cuts):
        # cut the training data
        tr_seqs = train_seqs_set[:n_train//(2**i)]
        # fit an SWI model to the training data
        swi_model = np.sum(tr_seqs, axis=0) + swi_prior
        swi_model = swi_model / np.sum(swi_model, axis=-1)[:, None]
        log_swi_model = np.log(swi_model)

        # Calculate the log likelihoods of testing sequence (to calculate perplexities later)
        probs_swi_test[r, i] = np.sum(test_seqs * log_swi_model)
        # Calculate the log likelihoods of sequences from p^\infty (to calculate KL later)
        probs_swi_infty[r, i] = np.sum(infty_sequences * log_swi_model)
        # Calculate mutation probabilities
        probs_swi_muts[r, i] = get_mut_probs(log_swi_model)
        for j, lag in enumerate(lags):
            # For each lag, preprocess training sequences and get lags
            (probs_bear_test[r, i, j], probs_bear_infty[r, i, j], probs_bear_train[r, i, j]
            ) = preprocess(tr_seqs, test_seqs, infty_sequences,
                           'sequences_r{}_c{}'.format(r, i), lag, max_lag, alphabet=alphabet_name,
                           bear_prior=np.array([bear_prior]), dump_seqs=(j==0))
            # now load preprocessed data into tensorflow data object
            data = get_kmers('sequences_r{}_c{}'.format(r, i), lag, max_lag, alphabet_name)
            # use the preprocessed data to sample mutaton probabilities from vanilla BEAR
            probs_bear_muts[r, i, j] = get_var_probs.get_bear_probs(None, wt_seq, vars_, 0,
                                                      mc_samples=mc_samples, vans=[bear_prior], lag=lag, alphabet=alphabet_name,
                                                      get_map=False, h=None, data=data)[:, 0, :]

# %% [markdown]
Before plotting, we need to select the MAP lag for BEAR

# %% [code]
lag_posterior = probs_bear_train - 4**(1.*lags)

probs_bear_test = probs_bear_test[np.arange(replicates)[:, None], np.arange(n_cuts)[None, :], np.argmax(lag_posterior, axis=-1)]
probs_bear_infty = probs_bear_infty[np.arange(replicates)[:, None], np.arange(n_cuts)[None, :], np.argmax(lag_posterior, axis=-1)]
probs_bear_muts = probs_bear_muts[np.arange(replicates)[:, None], np.arange(n_cuts)[None, :], np.argmax(lag_posterior, axis=-1)]

# %% [markdown]
Now we plot perplexities (fig 4C) to see that BEAR converges asymptotically to $p_0$ while SWI does not.

# %% [code]
import matplotlib.pyplot as plt
from matplotlib import cm
from matplotlib import patches

# %% [code]
swi_perplexity = np.exp(-probs_swi_test[:, ::-1]/(n_test*L))
bear_perplexity = np.exp(-probs_bear_test[:, ::-1]/(n_test)/L)


def plot_med(dat, color, label):
    plt.plot(np.median(dat, axis=0), color=color, label=label, linewidth=7)
    for i in range(len(dat)):
        plt.plot(dat[i], color=color, alpha=0.2, linewidth=3)
bear_colors = cm.viridis(np.linspace(0, 1, 4))

fig, ax = plt.subplots(1, figsize=[7, 7])
plt.xlabel("Training N", fontsize=30)
plt.xticks(np.arange(n_cuts)[::2], n_train//2**(np.arange(n_cuts)[::2][::-1]), fontsize=25)
for tick in ax.xaxis.get_major_ticks()[1::2]:
    tick.set_pad(25)
plt.ylabel("perplexity", fontsize=30)
plt.yticks(fontsize=25)
plot_med(swi_perplexity, "blue", "SWI")
plot_med(bear_perplexity, bear_colors[2], "BEAR $c=1$")
plt.legend(fontsize=25)
plt.tight_layout()
plt.ylim([1.0, 2.5])

# %% [markdown]
Now KL (Fig 4D) to see that while BEAR converges to $p_0$ it drifts from $p^\infty$

# %% [code]
swi_kl = (infty_ent - probs_swi_infty[:, ::-1]/n_infty)
bear_kl = (infty_ent - probs_bear_infty[:, ::-1]/n_infty)

fig, ax = plt.subplots(1, figsize=[7, 7])
plt.xlabel("Training N", fontsize=30)
plt.xticks(np.arange(n_cuts)[::2], n_train//2**(np.arange(n_cuts)[::2][::-1]), fontsize=25)
for tick in ax.xaxis.get_major_ticks()[1::2]:
    tick.set_pad(25)
plt.ylabel("$KL(p^{\infty}||p)$", fontsize=30)
plt.yticks(fontsize=25)
plot_med(swi_kl, "blue", "SWI")
plot_med(bear_kl, bear_colors[2], "BEAR $c=1$")
plt.legend(fontsize=25)
plt.ylim([0., 12])
plt.tight_layout()

# %% [markdown]
Now we also see that the SWI model, despite being trained on phylogenetically corellated data, can still capture the fitness landscape (S4).

# %% [code]
train_seqs_set = train_seqs[0]
# fit SWI to training data
tr_seqs = train_seqs_set[:n_train]
swi_model = np.sum(tr_seqs, axis=0) + swi_prior
swi_model = swi_model / np.sum(swi_model, axis=-1)[:, None]
log_swi_model = np.log(swi_model)

plt.figure(figsize=(8, 6))
plt.imshow(swi_model.T, cmap='Blues', vmin=0, vmax=1)
plt.title('fit SWI model distribution', fontsize=18)
plt.xlabel('position', fontsize=18)
plt.ylabel('nucleotide', fontsize=18)
cbar = plt.colorbar()
cbar.ax.tick_params(labelsize=18)

plt.figure(figsize=(8, 6))
plt.imshow(steady_state.T, cmap='Blues', vmin=0, vmax=1)
plt.title('steady state distribution', fontsize=18)
plt.xlabel('position', fontsize=18)
plt.ylabel('nucleotide', fontsize=18)
cbar = plt.colorbar()
cbar.ax.tick_params(labelsize=18)

# %% [markdown]
Finally we recreate figure 4B to show that BEAR, despite performing accurate density estimation, cannot accurately predict the probabilities of mutations.

# %% [code]
# set up a function that returns spearmans from mutation log probabilities
def ord_(mat):
    """Get back the ranks of the elements of an array"""
    return np.argsort(np.argsort(mat, axis=-1), axis=-1)
def norm(mat):
    """Centre and normalize an array"""
    return (mat - np.mean(mat, axis=-1)[..., None])/np.std(mat, axis=-1)[..., None]
def get_sp(mut_mat):
    """Calculate the spearman given an array of mutation probabilities"""
    return np.average(norm(ord_(mut_mat))*norm(ord_(true_mut_probs)), axis=-1)

# %% [code]
# functions to plot the data
def plot_mean(dat, color, label):
    plt.plot(np.mean(dat, axis=0), color=color, label=label, linewidth=7)
    for i in range(len(dat)):
        plt.plot(dat[i], color=color, alpha=0.2, linewidth=3)
        
def plot_graded_sps_scan(bear_perc, swi_sps, ylim=[-0.05, 1.05]):
    fig, ax = plt.subplots(1, figsize=[7, 7])
    plt.xlabel("Training N", fontsize=30)
    plt.xticks(np.arange(n_cuts)[::2], n_train//2**(np.arange(n_cuts)[::2][::-1]), fontsize=25)
    for tick in ax.xaxis.get_major_ticks()[1::2]:
        tick.set_pad(25)
    plt.ylabel("Spearman", fontsize=30)
    plt.yticks(fontsize=25)
    plt.ylim(ylim)
    
    colors = cm.viridis(np.linspace(0, 1, 4))
    num_cuts = len(bear_perc)

    #plot grid
    xlim = [-0.5, n_cuts-0.5]
    for i in np.arange(-0.0, 1.01, 0.1):
        plt.plot(xlim, [i, i], color='grey', alpha=0.7)
    plt.plot(xlim, [0, 0], color='black', alpha=1)
    plt.xlim(xlim)
            
    plot_mean(swi_sps, 'blue', 'SWI')
    plt.plot(bear_perc[:, 2], linewidth=5, color='black', label='BEAR')

    plt.errorbar(np.arange(num_cuts), bear_perc[:, 0],
                 yerr=[np.zeros(num_cuts), bear_perc[:, -1]-bear_perc[:, 0]], color=colors[2],
                 elinewidth=3, capsize=5, linewidth=0, markersize=0)
    for i in range(num_cuts):
        rect = patches.Rectangle((i-0.25, bear_perc[i, 1]),
                                 0.5, bear_perc[i, -2]-bear_perc[i, 1], linewidth=0, facecolor=colors[2])
        ax.add_patch(rect)

    plt.show()
    plt.close()

# %% [code]
swi_spearmans = get_sp(probs_swi_muts)[:, ::-1]
bear_spearmans = get_sp(np.transpose(probs_bear_muts[:, ::-1], [0, 1, 3, 2]))

for r in range(replicates):
    bear_perps = np.percentile(bear_spearmans[r], [2.5, 25, 50, 75, 97.5], axis=-1).T
    plot_graded_sps_scan(bear_perps, swi_spearmans[[r]], ylim=[0.49, 1.01])

# %% [code]
```
