## Supplemental code for "Non-identifiability and the Blessings of Misspecification in Models of Molecular Fitness": KingmanCoalescent.py

```
import numpy as np
from scipy.special import logsumexp
import scipy.stats as stats
import tensorflow_probability as tfp
from ete3 import Tree


def random_pair(lst):
    """Return random pair from a list, along with the remainder of the list."""
    indx = np.sort(np.random.choice(len(lst), 2, replace=False))[::-1]
    one = lst.pop(indx[0])
    two = lst.pop(indx[1])

    return one, two


class node:

    def __init__(self, t, A, B):

        self.t = t
        self.A = A
        self.B = B


def sample_coalescent(n):
    """Sample a genealogical tree from the Kingman coalescent."""
    # Make set of nodes.
    partition = [None]*n
    t = 0.
    for i in range(n-1):
        len_partition = len(partition)
        A, B = random_pair(partition)
        t -= stats.expon.rvs(scale=2/(len_partition*(len_partition-1)))
        new_node = node(t, A, B)
        partition.append(new_node)

    return partition


def OU_propagator(z0, delta_t, rate=1/2):
    """Propagate point z0 according to OU process."""
    loc = z0*np.exp(-rate*delta_t)
    scale = np.sqrt(1 - np.exp(-2*rate*delta_t)) / np.sqrt(2*rate)
    # Return parameters of normal distribution.
    return loc, scale


def sample_pou(coalescent, dim, rate=1/2):
    """Given a tree, sample from the phylogenetic OU process."""
    z_root = stats.norm.rvs(size=dim)
    xs = []

    # Setup recursion.
    def get_children(z0, t0, node):
        if node is not None:
            t1 = node.t
            # Propagate OU process to next node.
            loc, scale = OU_propagator(z0, t1-t0, rate=rate)
            z1 = stats.norm.rvs(loc=loc, scale=scale)
            # Fork.
            get_children(z1, t1, node.A)
            get_children(z1, t1, node.B)
        else:
            # Terminate recursion at leaf.
            loc, scale = OU_propagator(z0, -t0, rate=rate)
            x = stats.norm.rvs(loc=loc, scale=scale)
            xs.append(x)

    # Run recursion.
    get_children(z_root, np.finfo('float64').min, coalescent[0])

    return np.stack(xs)


def sample_SH_parameters(L, B, energy_scale):
    """Sample a fitness landscape for the Sella&Hirsh model."""
    h = (energy_scale * np.random.randn(L, B) /
         np.random.exponential(size=L)[:, None])
    return h


def compute_SH_dynamics(h, N, mu):
    """Solve S&H dynamics equations."""
    # Compute steady state distribution (Eqn. 7 in S+H)
    nu = 2 * (N - 1)
    steady_state = np.exp(-nu * h - logsumexp(-nu * h, axis=-1, keepdims=True))
    # Compute discrete-time transition probabilities.
    L, B = h.shape
    W = np.zeros((L, B, B))
    for b in range(B-1):
        for bp in range(b+1, B):
            h_delta = h[:, bp] - h[:, b]
            sign = np.sign(h_delta)
            W[:, b, bp] = mu * N * np.exp(
                                np.log(sign * np.expm1(2 * h_delta)) -
                                np.log(sign * np.expm1(2 * N * h_delta)))
            W[:, bp, b] = mu * N * np.exp(
                                np.log(- sign * np.expm1(- 2 * h_delta)) -
                                np.log(- sign * np.expm1(- 2 * N * h_delta)))
    for b in range(B):
        W[:, b, b] = 1 - np.sum(W[:, b, :], axis=-1)
    # Convert to continuous time dynamics
    A = np.transpose(W, axes=(0, 2, 1)) - np.eye(B)[None, :, :]
    # Solve to get full dynamics.
    Aeval, Aevec = np.linalg.eig(A)
    # Compute inverse to get eigenvector decomposition of basis vectors.
    Aevec_inv = np.linalg.inv(Aevec)

    return Aeval, Aevec, Aevec_inv, steady_state


def pSH_propagator(x, t, Aeval, Aevec, Aevec_inv):
    # Propagator for continuous time Markov dynamics.
    return np.einsum('ijk,ik->ij', Aevec,
                     np.exp(Aeval * t) * np.einsum('ijk,ik->ij', Aevec_inv, x))


def sample_pSH(coalescent, L, B, energy_scale=0.0001, pop_size=10000,
               rate=1.0):
    """
    Given a tree, sample from a phylogenetic Sella&Hirsh model with
    independent energies.
    L - gene length
    B - alphabet size (e.g. 4 or 20)
    energy_scale - scale for random energies (log fitnesses)
    pop_size - population size (N in Sella&Hirsh)
    rate - mutation rate (mu in Sella&Hirsh, assumed to be scalar)
    """
    # Sample fitness landscape.
    h = sample_SH_parameters(L, B, energy_scale)
    # Compute dynamics.
    Aeval, Aevec, Aevec_inv, steady_state = compute_SH_dynamics(
                                                    h, pop_size, rate)
    # Initialize with sample from steady_state.
    z_root = tfp.distributions.OneHotCategorical(
                probs=steady_state, dtype=np.float64).sample().numpy()
    xs = []

    # Setup recursion.
    def get_children(z0, t0, node):
        if node is not None:
            t1 = node.t
            # Propagate S&H continuous Markov process to next node.
            next_probs = pSH_propagator(z0, t1-t0, Aeval, Aevec, Aevec_inv)
            z1 = tfp.distributions.OneHotCategorical(
                        probs=next_probs, dtype=np.float64).sample().numpy()
            # Fork.
            get_children(z1, t1, node.A)
            get_children(z1, t1, node.B)
        else:
            # Terminate recursion at leaf.
            next_probs = pSH_propagator(z0, -t0, Aeval, Aevec, Aevec_inv)
            x = tfp.distributions.OneHotCategorical(
                        probs=next_probs, dtype=np.float64).sample().numpy()
            xs.append(x)

    # Run recursion.
    get_children(z_root, coalescent[0].t, coalescent[0])

    return np.stack(xs), h


def convert_to_ete(coalescent):
    """Convert the coalescent to ete3 format."""

    tree = Tree()
    seen_nodes = []

    def add_children(tree_node, coalescent_node):
        for branch in ['A', 'B']:
            new_coalescent_node = getattr(coalescent_node, branch)
            if new_coalescent_node is not None:
                seen_nodes.append(1)
                new_tree_node = tree_node.add_child(
                          name='{}'.format(sum(seen_nodes)),
                          dist=new_coalescent_node.t-coalescent_node.t)
                add_children(new_tree_node, new_coalescent_node)
            else:
                seen_nodes.append(1)
                tree_node.add_child(name='{}'.format(sum(seen_nodes)),
                                    dist=0.-coalescent_node.t)

    add_children(tree, coalescent[0])

    return tree
```
