## Supplemental code for "Non-identifiability and the Blessings of Misspecification in Models of Molecular Fitness": sequences_r0_c1_kmers_kmc_stderr_2_suf_2.txt

--- kmc ---

Info: Small k optimization on!

******* configuration for small k mode: *******
No. of input files           : 1
Output file name             : seqs/sequences_r0_c1_kmers_kmc_inter_2_suf_2.res
Input format                 : FASTQ

k-mer length                 : 2
Max. k-mer length            : 256
Min. count threshold         : 1
Max. count threshold         : 1000000000000
Max. counter value           : 1000000000000
Both strands                 : false
Input buffer size            : 33554432

No. of readers               : 1
No. of splitters             : 15

Max. mem. size               : 18000MB

Max. mem. for PMM (FASTQ)    :  3765MB
Part. mem. for PMM (FASTQ)   :    33MB
Max. mem. for PMM (reads)    :    23MB
Part. mem. for PMM (reads)   :     0MB
Max. mem. for PMM (b. reader):   402MB
Part. mem. for PMM (b. reader):   134MB


Stage 1: 0%
Stage 1: 100%
Warning: in 1 input file(s) there was not end of line character at EOF.
--- kmc dump ---
Warning: input database is already sorted. Each sort operation will be omitted

in1: 0% 
in1: 8% 
in1: 16% 
in1: 25% 
in1: 33% 
in1: 41% 
in1: 50% 
in1: 58% 
in1: 66% 
in1: 75% 
in1: 83% 
in1: 91% 
in1: 100% 
