## Supplemental code for "Non-identifiability and the Blessings of Misspecification in Models of Molecular Fitness": sequences_r0_c14_kmers_kmc_stderr_1_pre_6.txt

--- kmc ---

Info: Small k optimization on!

******* configuration for small k mode: *******
No. of input files           : 1
Output file name             : seqs/sequences_r0_c14_kmers_kmc_inter_1_pre_6.res
Input format                 : FASTQ

k-mer length                 : 6
Max. k-mer length            : 256
Min. count threshold         : 1
Max. count threshold         : 1000000000000
Max. counter value           : 1000000000000
Both strands                 : false
Input buffer size            : 33554432

No. of readers               : 1
No. of splitters             : 15

Max. mem. size               : 18000MB

Max. mem. for PMM (FASTQ)    :  3765MB
Part. mem. for PMM (FASTQ)   :    33MB
Max. mem. for PMM (reads)    :    23MB
Part. mem. for PMM (reads)   :     0MB
Max. mem. for PMM (b. reader):   402MB
Part. mem. for PMM (b. reader):   134MB


Stage 1: 0%
Stage 1: 100%
Warning: in 1 input file(s) there was not end of line character at EOF.
--- kmc dump ---
Warning: input database is already sorted. Each sort operation will be omitted

in1: 0% 
in1: 2% 
in1: 5% 
in1: 8% 
in1: 11% 
in1: 13% 
in1: 16% 
in1: 19% 
in1: 22% 
in1: 25% 
in1: 27% 
in1: 30% 
in1: 33% 
in1: 36% 
in1: 38% 
in1: 41% 
in1: 44% 
in1: 47% 
in1: 50% 
in1: 52% 
in1: 55% 
in1: 58% 
in1: 61% 
in1: 63% 
in1: 66% 
in1: 69% 
in1: 72% 
in1: 75% 
in1: 77% 
in1: 80% 
in1: 83% 
in1: 86% 
in1: 88% 
in1: 91% 
in1: 94% 
in1: 97% 
in1: 100% 
